# Loss of Nkd1 is epistatic to loss of Axin2 in regulating Wnt signaling

**DOI:** 10.1101/2023.10.30.563773

**Authors:** Ian Bell, Haider Khan, Nathan Stutt, Matthew Horn, Teesha Hydzik, Whitney Lum, Victoria Rea, Emma Clapham, Lisa Hoeg, Terence J. Van Raay

## Abstract

Wnt signaling is a crucial developmental pathway involved in early development as well as stem cell maintenance in adults and its misregulation leads to numerous diseases. Thus, understanding the regulation of this pathway becomes vitally important. Axin2 and Nkd1 are widely utilized negative feedback regulators in Wnt signaling where Axin2 functions to destabilize cytoplasmic β-catenin, and Nkd1 functions to inhibit the nuclear localization of β-catenin. Here, we set out to further understand how Axin2 and Nkd1 regulate Wnt signaling by creating *axin2*^-/-^, *nkd1*^-/-^ single mutants and *axin2*^-/-^;*nkd1*^-/-^ double mutant zebrafish using sgRNA/Cas9. All three Wnt regulator mutants were viable and had impaired heart looping, neuromast migration defects, and behavior abnormalities in common, but there were no signs of synergy in the *axin2*^-/-^;*nkd1*^-/-^ double mutants. Further, Wnt target gene expression by qRT-PCR, and RNA-seq analysis and protein expression by mass spectrometry demonstrated that the double *axin2*^-/-^;*nkd1*^-/-^ mutant resembled the *nkd1*^-/-^ phenotype demonstrating that Axin2 functions upstream of Nkd1 and that loss of Nkd1 is epistatic to the loss of Axin2. In support of this, the data further demonstrates that Axin2 uniquely alters the properties of β-catenin-dependent transcription having novel readouts of Wnt activity compared to *nkd1*^-/-^ or the *axin2*^-/-^;*nkd1*^-/-^ double mutant. We also tested the sensitivity of the Wnt regulator mutants to exacerbated Wnt signaling, where the single mutants displayed characteristic heightened Wnt sensitivity, resulting in an eyeless phenotype. Surprisingly, this phenotype was rescued in the double mutant, where we speculate that cross-talk between Wnt/β-catenin and Wnt/Planar Cell Polarity pathways could lead to altered Wnt signaling in some scenarios. Collectively, the data emphasizes both the commonality and the complexity in the feedback regulation of Wnt signaling.

## Introduction

Canonical Wnt signaling is important in regulating many events during early development such as initiating the dorsal-ventral axis, maintaining the midbrain-hindbrain barrier, and regulating stem cell differentiation and maintenance^1–3^. Active canonical Wnt signaling has been found to play a crucial role in the development of a majority of the major organs, including the kidneys, liver, and colon^2,4,5^. Dysregulation of Wnt signaling, primarily due to mutations causing the pathway to become hyperactive, have been linked to many diseases including cancer, diabetes, bone density defects and neurological disorders to name a few^6–11^.

When Wnt signaling is off, cytoplasmic β-catenin is unstable due to a multi-protein complex termed the destruction complex^12,13^. The destruction complex is composed of Glycogen Synthase Kinase-3β (Gsk3β), Adenomatous Polyposis Coli (Apc), Axin, and Casein Kinase 1α (Ck1α), and functions to phosphorylate and ultimately destabilize cytoplasmic β-catenin^12,13^. Apc and Axin act as scaffolding proteins in the assembly of the destruction complex, whereas Gsk3β and Ck1α are kinases that phosphorylate β-catenin leading to its polyubiquitination and degradation by the proteasome^12,13^. At the transcriptional level, Wnt target genes remain silent due to transcriptional repressors, such as Groucho, binding to Tcf/Lef transcription factors^14^.

When Wnt signaling is on, a Wnt ligand binds its receptors Frizzled and Lipoprotein Receptor-Related Protein 5 and 6 (Lrp5/6) which recruits the cytoplasmic protein Dishevelled (Dvl) and the destruction complex to the membrane, resulting in the inactivation of the destruction complex^13^. Once this complex is deactivated, cytoplasmic levels of β-catenin stabilize allowing for the translocalization of β-catenin to the nucleus where it interacts with various coactivators to transcriptionally activate Wnt target genes^15^. Two of these Wnt target genes are *axin2* and *nkd1*, which function as negative feedback regulators for the pathway^16,17^.

Nkd1 is an antagonist in both the canonical and non-canonical Wnt signaling pathways^18–20^. At the N-terminal end of Nkd1 is an N-terminal myristoylation sequence which acts as a reversible membrane localization switch. The Nkd1 myristoylation and its localization to the membrane is necessary for its activity in both canonical and non-canonical Wnt signaling^21,22^. There is also an EF-hand in Nkd1 which is most similar to the Recoverin protein EF-Hand^23^. Recoverin is a myristoyl switch protein that changes its membrane localization upon calcium binding to the EF-Hand^24^. While calcium was found not to bind to the Drosophila EF-Hand of Nkd, it is not clear if calcium is required for vertebrate Nkd1 function^25^. Importantly, the EF-Hand of Drosophila is unique from other arthropods and vertebrate Nkd EF-Hands suggesting it has derived a novel function^19^. Nonetheless, the EF-Hand of Nkd1 is important for its ability to antagonize Wnt signaling as the EF-hand has been found to interact with several proteins including Dvl, β-catenin, and p62^22,23,25,26^. Furthermore, Nkd1 interaction with Dvl, via the EF-hand, is also important for its function in non-canonical Wnt signaling as Nkd1 constructs containing mutations in conserved residues in the EF-hand, are still able to rescue a Wnt8 induced eyeless phenotype in zebrafish but have altered non-canonical Wnt signaling^19^. There are other conserved domains including a homology domain that is also speculated to bind Dvl and a C-terminal poly histidine tail that is involved with its interaction with Axin and Axin2^22,26,27^. The model put forth by Gammons et al suggests that the poly-histidine tail is required for Nkd1’s aggregation that is dependent on its interaction with Dvl^26^. The aggregation of Nkd1 ensures that it only functions after sustained Wnt signaling^26^. Importantly, Nkd1’s function and interaction with β-catenin is Wnt ligand dependent, as Nkd1 was unable to inhibit Wnt signaling induced by ligand-independent mechanisms or in the absence of its myristoylation sequence^20,22,28^. Further, our model suggests that Nkd1 functions downstream of Dvl to inhibit the nuclear accumulation of β-catenin^22^.

Loss of Nkd1 in Drosophila results in a naked cuticle phenotype in the larva and in wing patterning defects later in development, both of which demonstrate that Nkd functions to inhibit canonical Wnt signaling^29,30^. In mice, loss of Nkd1 and/or Nkd2 has subtle defects in cranial bone morphology, slightly reduced litter sizes and effects on spermatogenesis, but are otherwise viable^31,32^. In zebrafish, knockdown of Nkd1 results in exacerbated Wnt signaling^20^. Finally, misexpression of Nkd’s in multiple systems further demonstrates that Nkd’s act as Wnt signaling antagonists^19,20,22,23,28,33^. Curiously, loss of Nkd1 in cell culture experiments has been shown to attenuate canonical Wnt signaling, suggesting that Nkd1 may function to promote Wnt signaling under certain conditions^26,27^.

Axin1 and Axin2 serve a similar function acting as scaffolding proteins in the assembly of the destruction complex. They have unique expression patterns with *axin1* being more ubiquitously expressed whereas *axin2* expression is more restricted ^34–36^. Furthermore, Axin1 appears to be a more potent inhibitor of canonical Wnt signaling as overexpression of Axin1 reduces β-catenin protein levels more than overexpression of Axin2 as seen by immunocytochemistry in SW480 cells^17^. However, when Wnt signaling is activated, Axin2 is a more potent antagonist for the pathway as knocking down Axin2, but not Axin1, increased dephosphorylated levels of β-catenin in the presence of the Wnt ligand revealing that Axin1 cannot compensate for the loss of Axin2^17^. Furthermore, both Axin1 and Axin2 have RGS and DIX domains which are important in facilitating polymerization, however, the domains are not identical to one another, allowing the proteins to have unique activity^37,38^. Mutating the DIX domain of both Axin1 and Axin2 inhibits their ability to self polymerize and reduces their ability to antagonize Wnt signaling^38^. Interestingly, the DIX domain of Axin1 is more sensitive to Dvl2 inhibition as chimeric Axin1 proteins, swapping in the DIX domain of Axin2, are no longer inhibited by Dvl2^17^. Also, an independent study found that Axin1, but not Axin2, binds to Dvls^27^. The unique response to Dvl proteins could allow Axin1 and Axin2 to have separate activity when the Wnt pathway is active. The differential importance for Axin1 and Axin2 during development is highlighted in knockout mouse models as loss of Axin1 is lethal whereas loss of Axin2 has no obvious phenotype during early embryonic development^39,40^. Later in development, Axin2 knockout mice have a small head phenotype, delayed heart valve maturation, and thickened aortic valve but are viable and fertile^41,42^.

In vitro, it has been shown that Nkd1 interacts with both Axin1 and Axin2^27^. Furthermore, Nkd1 forms a ternary complex with Dvl2 and Axin1. Axin1 only interacted with a high molecular weight aggregate form of Nkd1 with the interaction being abolished by mutating the C-terminal histidine cluster of Nkd1^26^. Nkd1 interaction with Axin1 could be important for its function as knocking down Axin1 in HEK293T cells reduced Nkd1’s ability to antagonize Wnt3a activity^27^. Furthermore, HEK293T cells with the C-terminal histidine cluster mutated in Nkd1 had reduced Wnt3a topflash activity revealing a possible positive regulatory function for Nkd1^26^. Collectively, these results suggest that Axins and Nkd1 proteins function at the level of β-catenin to antagonize Wnt signaling.

Current research has revealed that Wnt target gene activation is dynamic depending on the coactivators that interact with β-catenin^43^. In the development of colorectal cancer, there is the “just right” hypothesis for Wnt signaling to provide the “right” conditions for tumour growth^44^. This hypothesis was established by analyzing second hit *APC* mutations found in colorectal adenomas from familial adenomatous polyposis patients with a majority of second hits being point mutations which would disrupt β-catenin binding rather than completely abolishing the proteins activity^44^. Through these experiments we are beginning to understand that different levels of Wnt signaling have different consequences, however, investigation into understanding this concept can be challenging.

Nkd1 and Axin2 are both Wnt target genes, and both interact with each other as well β-catenin to inhibit Wnt signaling. Axin2 function is ligand independent and acts to degrade β-catenin while Nkd1 activity is ligand dependent and acts to prevent β-catenin’s nuclear accumulation. The interaction between Axins and Nkds is curious given their unique modes of action and it is unclear how they would function together to inhibit Wnt signaling. Alternatively, it could be that the interaction is merely a mechanism to aggregate Wnt signaling components together to nucleate inhibitors^45^. Based on available information, we speculate that these two regulators function independently of one another to provide differential regulation of the pathway, not just on or off as we now know that different levels of β-catenin have different consequences on gene transcription and ultimately cell function. Thus, we hypothesized that compared to both single mutants, the combined loss of Axin2 and Nkd1 would result in a synergistic phenotype, resulting in hyperactivated or hypersensitized Wnt signaling. Counter to our hypothesis, we found that Axin2 and Nkd1 function in a classic epistatic scenario, where Nkd1 functions downstream of Axin2 and loss of Nkd1 is epistatic to loss of Axin2.

## Results

### Knocking out Axin2 and Nkd1

The Axin2 and Nkd1 knockout lines were created by injecting sgRNA/Cas9 with their respective sgRNAs at the one cell stage and grown to adulthood (P0). These were back crossed into their respective genetic lines and F1 founders were tail clipped and genotyped via indel detection by amplicon analysis followed by sequencing. We evaluated several mutants for each gene, all of which showed similar phenotypes (not shown) and thus we only focused on one genotype for each of *axin2* and *nkd1*. The *axin2*^-/-^ zebrafish were made in the tail long background (TL) whereas the *nkd1*^-/-^ zebrafish were generated in the Tubingen (TU) background. The *axin2*^-/-^ mutant has a 12-nucleotide deletion with a 1 nucleotide insertion in exon 2 leading to a frameshift mutation and premature stop codon (Supplemental Fig 1). The *nkd1*^-/-^ mutation is a 16-nucleotide deletion in exon 3 leading to a frameshift mutation and premature stop codon (Supplemental Fig 2). The *axin2*^-/-^;*nkd1*^-/-^ zebrafish were created by crossing *axin2*^+/-^ zebrafish with *nkd1*^-/-^ to create *axin2*^+/-^;*nkd1*^+/-^. The *axin2*^+/-^;*nkd1*^+/-^ zebrafish were then crossed to *nkd1*^-/-^ to create *axin2*^+/-^;*nkd1*^-/-^, in which *axin2*^+/-^;*nkd1*^-/-^ zebrafish were incrossed to make *axin2*^-/-^;*nkd1*^-/-^.

For reference, in the figure legends, capital “N” refers to the number of biological replicates, whereas lowercase “n” refers to the total number of observations. In addition, all mutants discussed below are maternal-zygotic mutants unless otherwise specified.

### Wnt regulator mutants have developmental defects

Our first objective was to determine if the different mutants had developmental defects and to report on those that we consistently observe. While some of the experiments reported below have only 1 biological replicate, all have been observed multiple times and in different genetic backgrounds. We report only on the ones where we have included all the relevant controls.

The Wnt regulator mutants have no overt phenotype, appear phenotypically wild type until early adult hood (see below), and are viable. Upon close examination it was demonstrated that the mutants have heart looping defects (Fig. 1A). Specifically, at 2 days post fertilization (dpf), wild type larvae had D looped hearts 90.6% of the time, whereas, the *axin2*^-/-^, *nkd1*^-/-^, and *axin2*^-/-^;*nkd1*^-/-^ had D looped hearts 41.3%, 53.8%, and 26.9% of the time, respectively. In each of the Wnt regulator mutants there was an increase in proportion of D incomplete looped, L looped, L incomplete looped, and unlooped hearts when compared to wild type. Since heart looping phenotypes can be variable based on the genetic background, we injected wild type embryos with sgRNA/Cas9 targeting both Axin2 and Nkd1 and observed a similar proportion of heart looping phenotypes compared to the knockout lines (Fig. 1B). These results support research on Wnt signaling in heart development where *apc* mutant zebrafish also develop with unlooped hearts, although at a much greater proportion^46^. Furthermore, past research has demonstrated that knock down of Nkd1 with morpholinos impairs heart development by inducing a heart jogging defect^33^.

**Figure 1.**
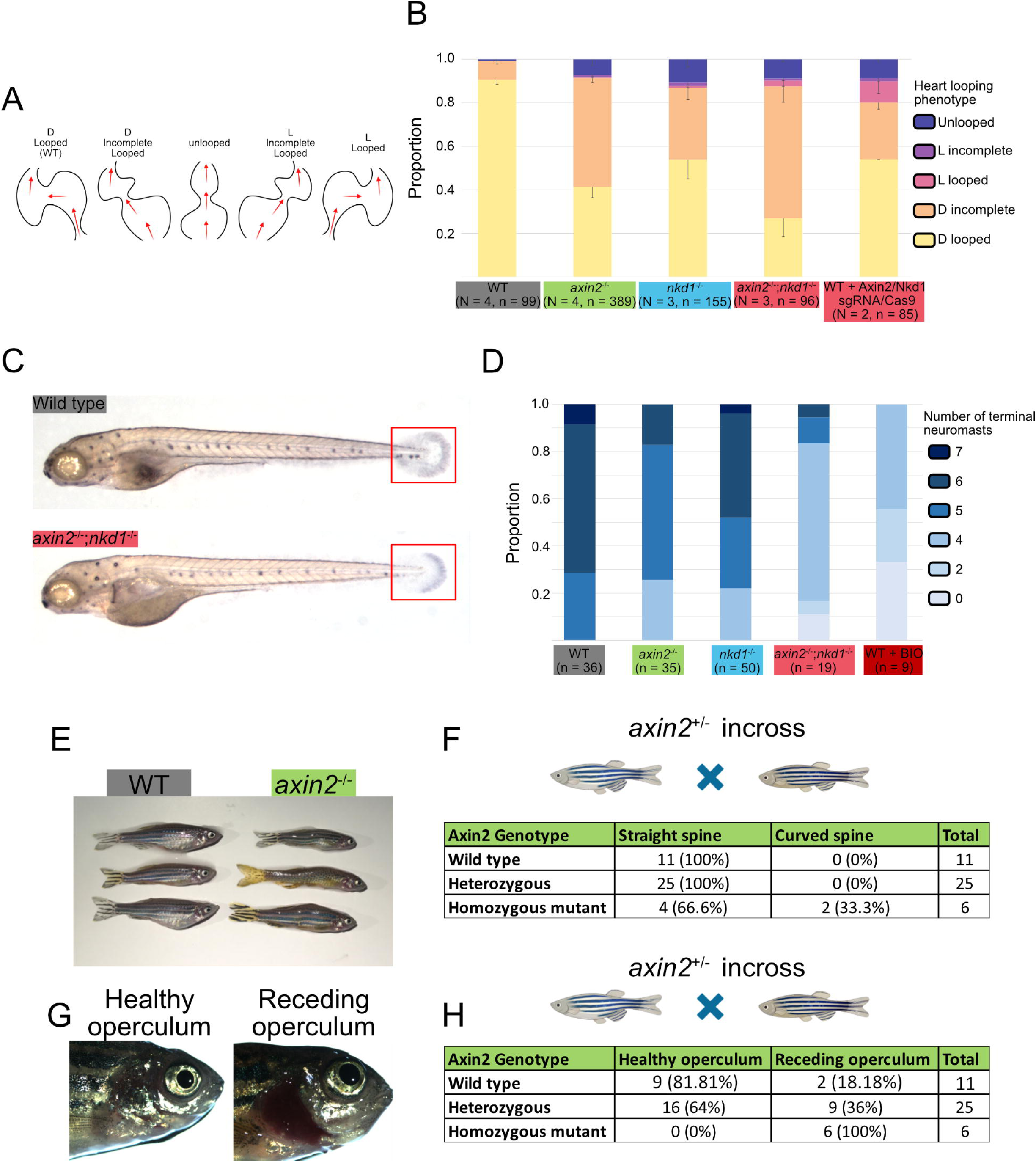
Wnt regulator mutants have developmental defects. (A) 2dpf Zebrafish embryos were observed for heart looping phenotypes and quantified in (B). (C) Zebrafish were treated with PTU from 1dpf - 5dpf and then fixed at 5dpf for neuromast staining with terminal neuromasts highlighted by the red box and quantified in (D). (E) Maternal-zygotic adult *axin2*^-/-^ zebrafish occasionally develop a curved spine phenotype. (F) Adults from a heterozygous incross were phenotyped for the curved spine followed by genotyping. (G) Maternal-zygotic adult *axin2*^-/-^ zebrafish often have a receding operculum phenotype. (H) Adults from a heterozygous incross were phenotyped for the receding operculum followed by genotyping. Error bars represent SEM.

We also examined other known Wnt dependent events for signs of activated Wnt signaling. Wnt signaling is necessary for neuromast formation and migration where it works alongside FGF signaling for proper patterning^47–49^. Hyperactivation of Wnt signaling has previously been shown to reduce neuromast migration in zebrafish which lead us to investigate if neuromast migration is impaired in the Wnt regulator mutants (Fig. 1C)^47^. At 5dpf, wild type zebrafish had 5-7 terminal neuromast, whereas, *axin2*^-/-^, *nkd1*^-/-^, and *axin2*^-/-^;*nkd1*^-/-^ double mutants had 4-6, 4-7, and 0-6 terminal neuromast, respectively. To confirm that hyperactivation of Wnt causes reduced neuromast migration, wild type embryos were treated with a small molecule Wnt activator, 6-bromoindirubin-3’-oxime (BIO), which caused larvae to have 0-4 terminal neuromasts at 5dpf (Fig. 1D). Total number of neuromasts for all Wnt regulator mutants did not change (data not shown) suggesting that the migration of the neuromasts are perturbed, consistent with published reports^47^.

We also noticed a curved spine phenotype in the *axin2*^-/-^ mutants. An incross of *axin2*^+/-^ mutants resulted in 42 adults, of which, 2/6 *axin2*^-/-^, 0/25 *axin2*^+/-^, and 0/11 *axin2*^+/+^ developed with a curved spine phenotype (Fig. 1E-F). An incross of *axin2*^-/-^ mutants produced a similar frequency of phenotypes (∼20-50% of homozygous mutants), suggesting that there is no maternal contribution. The *axin2*^-/-^;*nkd1*^-/-^ zebrafish also develop a curved spine, albeit with less frequency, whereas *nkd1*^-/-^ single mutants do not (data not shown). This suggests that the genetic background (TL in the *axin2*^-/-^ mutants and predominantly TU in the *axin2*^-/-^;*nkd1*^-/-^ mutants) does not have a significant role. This phenotype is not surprising given Wnt signaling is important in bone formation where both hyperactivation and hypoactivation of the pathway leads to skeletal deformities^40,50,51^. Furthermore, Axin2 knockout mice have axial skeletal defects and a small head phenotype^40^.

Further, the *axin2*^-/-^ adult zebrafish often develop with receding opercula (Fig. 1G-H). In contrast to the low to moderate penetrance of the curved spine, *axin2*^-/-^ mutants from an incross of *axin2*^+/-^ had complete penetrance (6/6) of receding opercula in the one cross that we performed whereas *axin2*^+/-^ and axin2^+/+^ had 9/25 and 2/11 with receding opercula, respectively, suggesting that there may be a dosage effect (Fig 1H). Operculum malformation is common in stressed or old age zebrafish, however, *axin2*^-/-^ mutants consistently develop with a receding operculum at an early age (∼5 months). The *axin2*^-/-^;*nkd1*^-/-^ often have receding opercula at a frequency seen in the *axin2*^-/-^ mutants, whereas, *nkd1*^-/-^ mutants do not (data not shown).

The varying degree in the number of terminal neuromasts suggests that the Wnt regulator mutant larva might have altered responses to stimulation. Therefore, we assessed 5dpf Wnt regulator mutants for swimming behavior and tap response. The larvae were recorded for 1.5 hours under dark conditions using a 24 well plate and assayed for thigmotaxis. We discovered that the wild type larvae had a greater thigmotactic behavior when compared to the Wnt regulator mutants. Furthermore, while wild type larvae spent 82.6% of their time along the edge of the well, *axin*2^-/-^, *nkd1*^-/-^, and *axin2*^-/-^;*nkd1*^-/-^ mutants spent 70.6%, 70.9%, and 75.4%, of their time along the edge, respectively. Startle response was measured for 30 taps over a period of 30 seconds with the first three taps in the analysis showing the greatest difference in response when comparing the Wnt regulator mutants to wild type larvae. The *axin2*^-/-^ larvae had a mild reduction in startle response whereas the *nkd1*^-/-^ and *axin2*^-/-^;*nkd1*^-/-^ larvae had a severe reduction in startle response when compared to wild type (Supplemental Fig. 3).

### Wnt regulator mutants have differential sensitivities to exogenous Wnt8

While Axin2 and Nkd1 are both negative feedback regulators of Wnt signaling acting at the level of β-catenin, we have previously demonstrated that exogenous Nkd1 is only active in the presence of ligand activated Wnt signaling^28^. In contrast, Axin2 does not require the Wnt ligand for its activation^28^. Therefore, we wanted to assess the response of the regulator mutants to overactive Wnt signaling. Overexpression of *wnt8* leads to an eyeless phenotype that is easily visible at 1dpf (Fig. 2A-B). Given that both Axin2 and Nkd1 are negative feedback regulators, we hypothesized that loss of these regulators would exacerbate the eyeless phenotype induced by Wnt8 overexpression. Injection of a low dose of *wnt8* where only 4.7% of embryos are eyeless in wild type, increased to 33.8% and 31.6% in *axin2*^-/-^ and *nkd1*^-/-^ mutants respectively, supporting our hypothesis (Fig. 2C). We further expected the *axin2*^-/-^;*nkd1*^-/-^ double mutants to be even more sensitive than either of the single mutants but surprisingly, only 4.6% of *axin2*^-/-^;*nkd1*^-/-^ larvae were eyeless, suggesting the loss of one regulator rescues the effect of the loss of the other (Fig. 2C). Furthermore, *nkd1*^-/-^ injected mutants had a large proportion of embryos develop with a short-twisted axis when *wnt8* was overexpressed, which was rarely seen in either *axin2*^-/-^ single or *axin2*^-/-^;*nkd1*^-/-^ double mutants (Fig. 2D-F). This suggests that loss of Axin2 can rescue loss of Nkd1 in the short- twisted axis phenotype, but it is unknown if this is the case in the eyeless phenotype.

**Figure 2.**
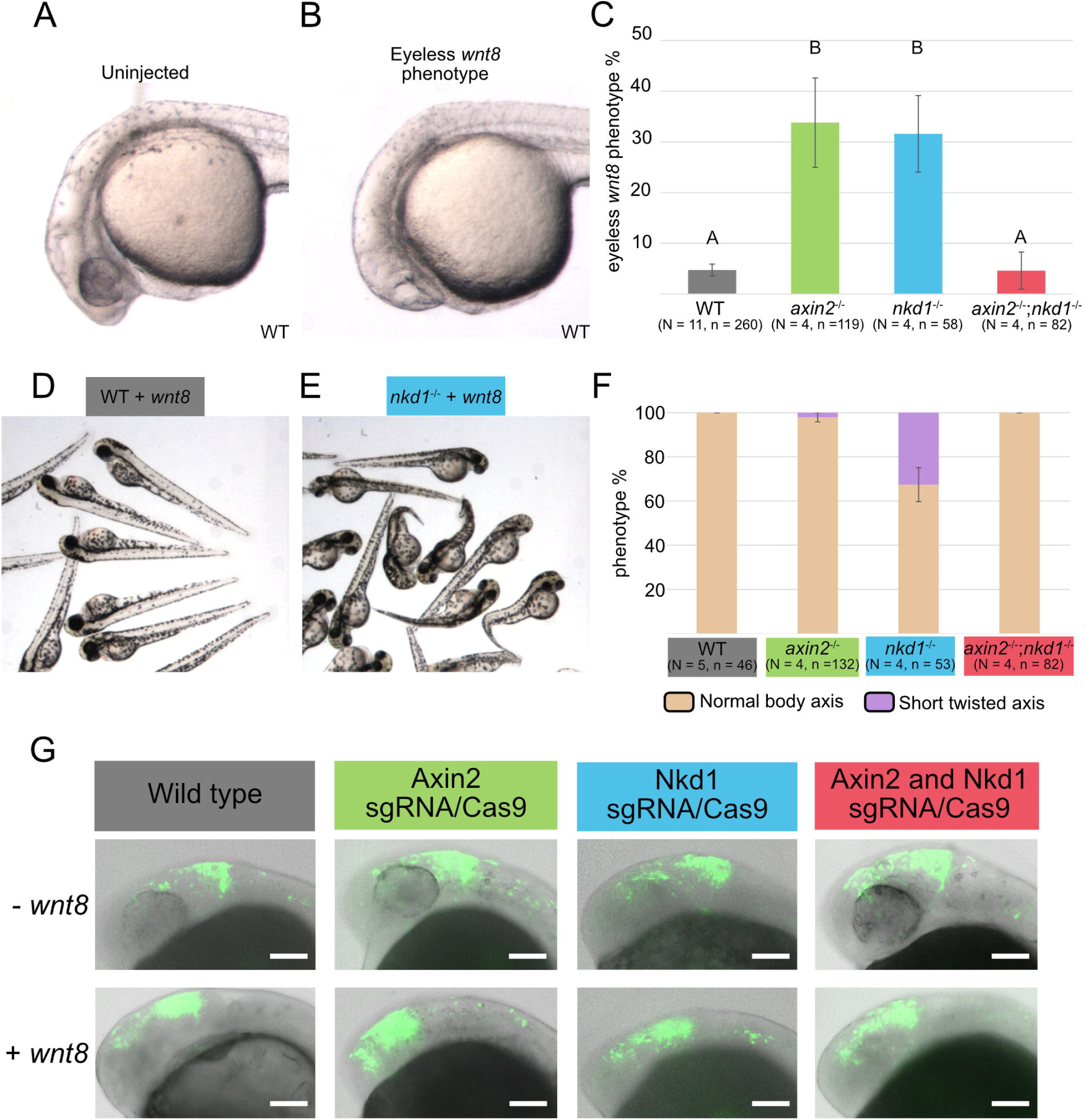
Wnt regulator mutants have differential sensitivities to exogenous Wnt8. (A-C) Overexpression of 100 pg of *wnt8* in wild type embryos results in the classic Wnt gain of function eyeless phenotype in approximately 5% of embryos, but more than 30% in *axin2*^-/-^ and *nkd1*^-/-^ mutants. Small eyes were considered to have eyes. In contrast, the *axin2*^-/-^;*nkd1*^-/-^ resembles the wild type phenotype (C). (D-F) Analysis of body axis development revealed that only the *nkd1*^-/-^ mutants develop a short-twisted axis with exogenous *wnt8* in about 30% of the embryos. (G) The Wnt reporter line was used to assess Wnt activity at the midbrain-hindbrain boundary at 1dpf when either Axin2 and/or Nkd1 was knocked down using sgRNA/Cas9. In each of the sgRNA/Cas9 injections there was an expansion of Wnt activity in the forebrain when compared to wild type embryos. Single plane brightfield images were overlaid with composite GFP images and as such the eye is not visible in all images. Error bars represent SEM, different letters above the bar signify significance with a p-value < 0.01 using a one-way ANOVA.

To further explore the effect of knocking out Axin2 and/or Nkd1 on Wnt activity we utilized the Wnt reporter zebrafish line (7XTCF:GFP)^52^. We injected *axin2* and/or *nkd1* sgRNAs along with Cas9 protein into one-cell stage transgenic embryos derived from the Wnt reporter line (Crispants). Wnt activity via GFP expression in the brain was then analyzed at 1dpf. Uninjected wild type embryos predominantly have GFP expression at the midbrain-hindbrain boundary with some activity being present in the forebrain (Fig. 2G). Knock down of Axin2 caused an anterior-ward expansion in GFP expression in the forebrain, whereas knock down of Nkd1 only caused a mild increase in GFP expression in the forebrain. When both Axin2 and Nkd1 were knocked down there was a similar expansion in GFP expression as seen in the *axin2* crispants alone, suggesting the double crispant is more similar to the *axin2* crispant but the variability in the data precludes statistical testing. To further explore this, we overexpressed *wnt8,* which caused *axin2* crispants to have a major anterior expansion in GFP expression which was milder in the *nkd1* or *axin2*;*nkd1* crispant embryos (Fig. 2G). Importantly, the eyeless phenotype induced in this transgenic line mimicked the knockouts where the single crispants had increased sensitivity while the double crispants showed the same insensitivity to exogenous Wnt8 (data not shown).

Thus far, loss of either Axin2 or Nkd1 produces results that are consistent with their roles as negative feedback regulators; however, things get confusing when both Axin2 and Nkd1 are simultaneously knocked out, especially in response to exogenous Wnt8. Therefore, we wanted to test how these mutants responded when Wnt signaling was overactivated in a ligand independent manner. To accomplish this, we used the small molecule BIO, which inhibits Gsk3β activity (Fig. 3A,B)^53^. Wild type embryos treated with 0.5µM of BIO reduces the size of the eye approximately 45.7% (±19.9%) (Fig. 3C). Using the same dose of BIO, *axin2*^-/-^, *nkd1*^-/-^, and *axin2*^-/-^;*nkd1*^-/-^ double mutant eyes were reduced by 69.5% (±21.6%), 96.1% (±3.9%), and 89.4% (±0.42%), respectively (Fig. 3C, supplemental Fig. 4). To ensure that the sensitivity to BIO was not a consequence of the different genetic backgrounds we created *axin2*^-/-^ mutants from an *axin2*^+/-^ incross. Here, *axin2*^-/-^ embryos treated with BIO had an increase in the eyeless phenotype when compared to embryos with at least one copy of wild type Axin2 from the same cross demonstrating that the sensitivity is specific for loss of Axin2 (Supplemental Fig. 5). In stark contrast to the effect of ectopic Wnt8, the *axin2*^-/-^;*nkd1*^-/-^ double mutant was highly sensitive to BIO. This suggests that a ligand, or receptor proximal, dependent event is somehow inhibiting Wnt/β-catenin signaling in the absence of two intracellular negative feedback regulators during early gastrulation when canonical Wnt signaling is patterning the hindbrain^54^.

**Figure 3.**
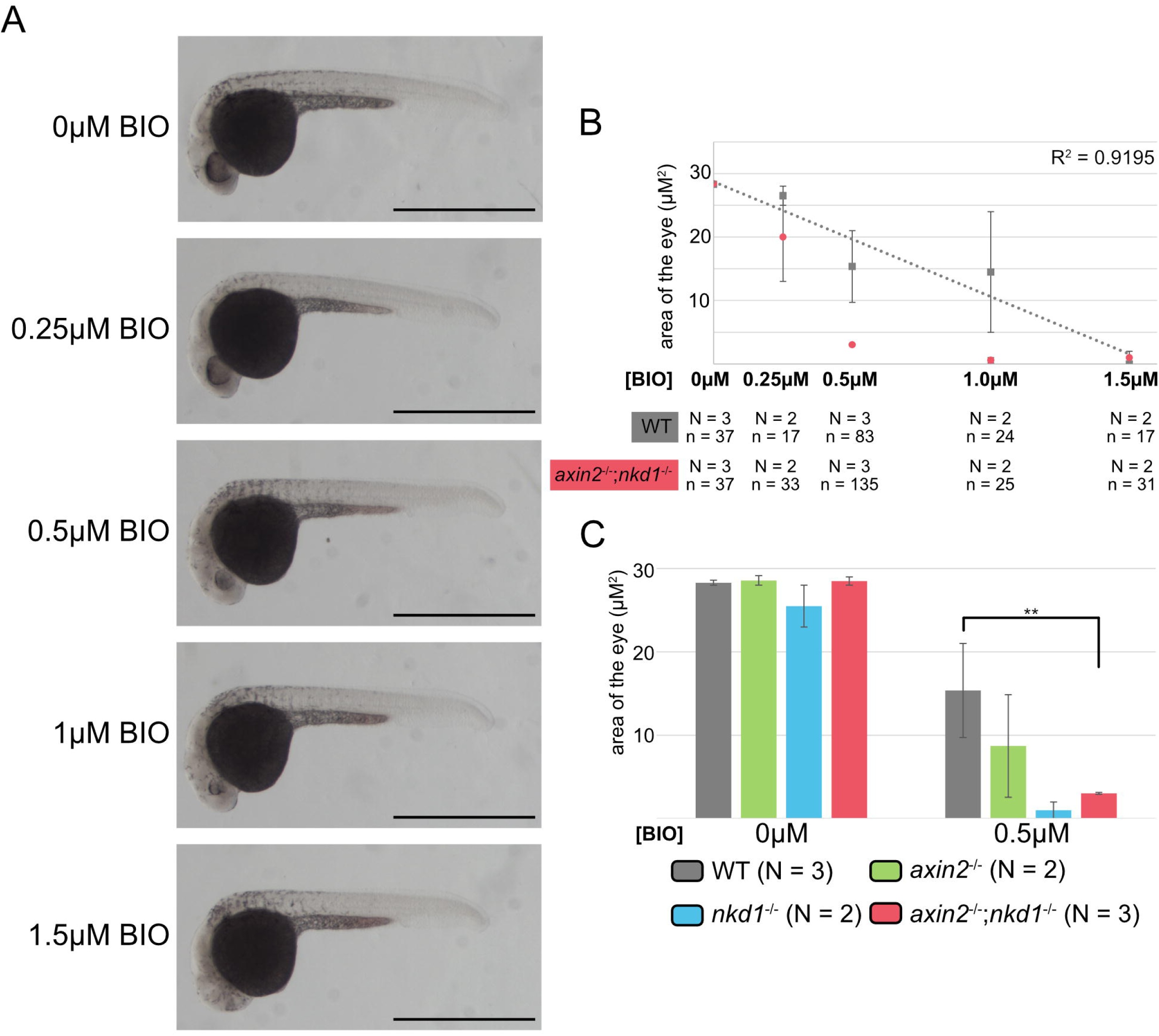
*axin2*^-/-^;*nkd1*^-/-^ mutants are sensitive to activation at the destruction complex level. (A) Embryos were treated with varying concentrations of BIO from 4 – 30 hpf with representative images taken of the wild type embryos. (B) In wild type embryos, the area of the eye has a decreasing linear association with the concentration of BIO with all embryos being eyeless at 1.5µM. In contrast, the majority of the *axin2*^-/-^;*nkd1*^-/-^ mutant embryos were eyeless at 0.5µM of BIO. (C) The *axin2*^-/-^;*nkd1*^-/-^ mutant has no change in the size of eye in the control treatment but has significantly smaller eyes at 0.5µM of BIO when compared to wild type revealing that the *axin2*^-/-^;*nkd1*^-/-^ mutants are sensitive to BIO. Measurements were performed on imageJ and a t-test was used with ** = p-value < 0.01. All error bars represent SEM. No statistics were performed on the *axin2*^-/-^ and *nkd1*^-/-^ single mutants.

### Wnt regulator mutants have differential effects on Wnt target genes

The data so far suggests that regulation of Wnt by negative feedback regulators is a complex process. To further explore the effect of Wnt8 and determine if we could separate out how these two regulators function to restrict Wnt signaling, we analyzed the expression of several well characterized Wnt target genes. At 30% and 50% epiboly *axin2* expression is restricted to the ventral lateral domain of the developing zebrafish embryo. Overexpression of Axin1 abolished *axin2* expression at 30% and 50% epiboly, whereas hyperactivating Wnt signaling using a stable form of β-catenin (ΔN-β-catenin) causes ectopic *axin2* expression (Supplemental Fig. 6). Combined with published results, this confirms that Axin2, along with Nkd1 and Sp5a are excellent markers for active Wnt signaling during pre-gastrulation development^55–58^.

We first used qRT-PCR to quantify the expression of the Wnt target genes *axin2*, *nkd1*, and *sp5a* at 30% and 50% epiboly (when zygotic Wnt signaling is high) with or without overexpression of *wnt8* in the different genetic backgrounds (Fig 4)^20,59,60^. We did not observe a decrease in endogenous *axin2* or *nkd1* expression in any of the Wnt regulator mutants, suggesting that the mutations in *axin2* and *nkd1* are not inducing nonsense mediated decay. In the *axin2*^-/-^ mutants, there was a predictable, albeit slight, increase in all three target genes that was exacerbated in the presence of exogenous Wnt8 at both 30% and 50% epiboly. Curiously, only the expression of *axin2* in the *axin2*^-/-^ mutants was significant at 30% epiboly (Fig. 4A,B). Conversely, the *nkd1* ^-/-^ mutants displayed a somewhat different pattern, as there was no substantive increase in either *axin2* or *nkd1* expression at 30% with or without exogenous Wnt8 when compared to controls (Fig. 4C). At 50%, the expression of *axin2* and *nkd1* in *nkd1*^-/-^ was similar to the *axin2*^-/-^ mutant in that there was a slight, but non-significant increase in their expression that was moderately enhanced by exogenous Wnt8 (Fig. 4D). Interestingly and curiously, *sp5a* expression in uninjected *nkd1*^-/-^ mutants was not significantly impacted but skyrocketed in the presence of exogeneous Wnt8 at 30% epiboly (Fig. 4C). Furthermore, at 50% epiboly, uninjected *nkd1*^-/-^ mutants had a significant increase in *sp5a* expression that was further exacerbated by the addition of exogenous Wnt8 (Fig. 4D). Importantly, the *axin2*^-/-^;*nkd1*^-/-^ mutant closely resemble the *nkd1*^-/-^ mutants with respect to all three genes (Fig. 4E). This strongly suggests that each single mutant has a unique consequence on the activation of Wnt target genes and that loss of Nkd1 might be epistatic to the loss of Axin2.

**Figure 4.**
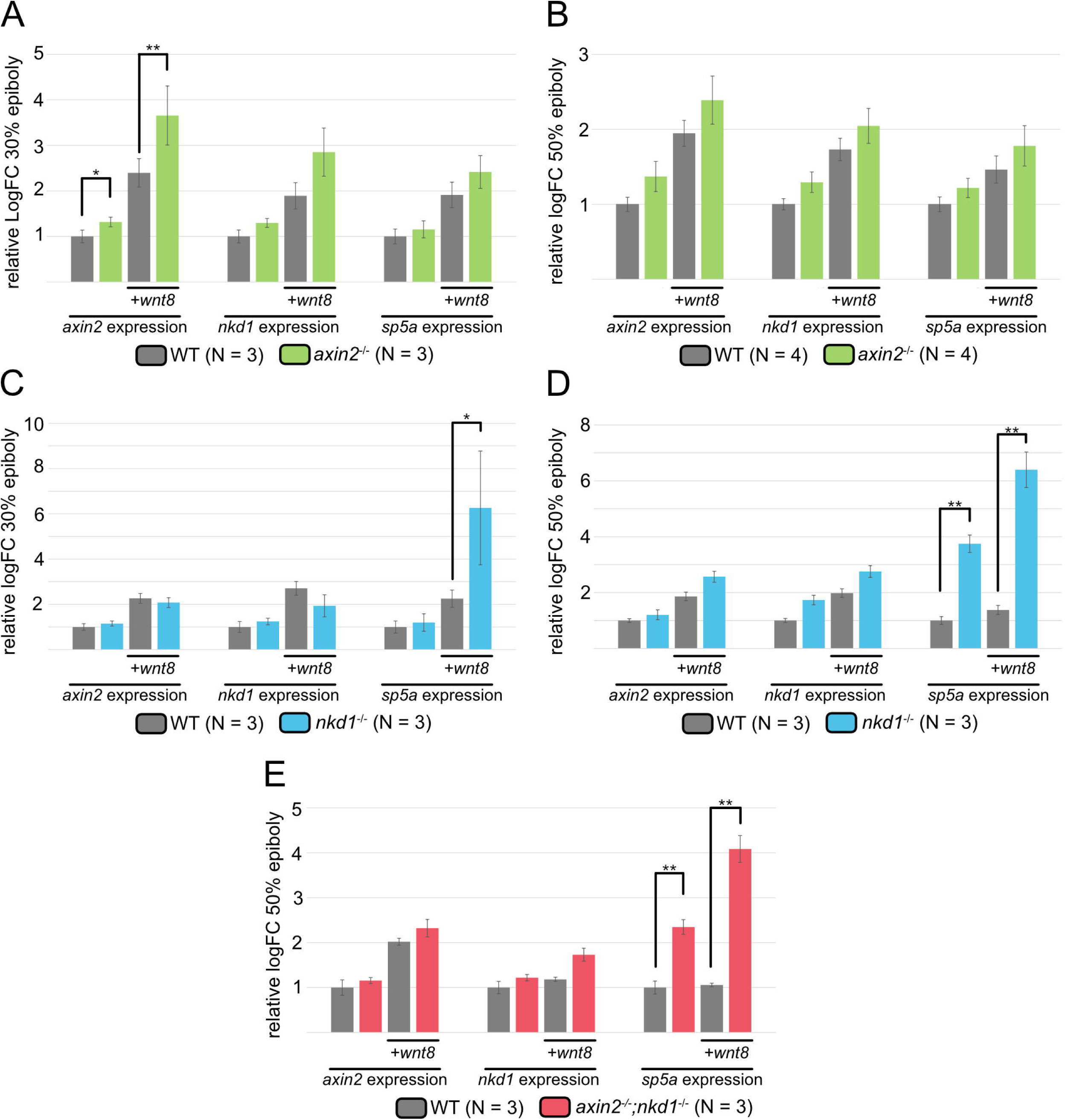
Wnt regulator mutants have differential effects on Wnt target genes. qRT-PCR was performed at 30% (A,C) and 50% (B, D, E) epiboly for the Wnt target genes *axin2*, *nkd1*, and *sp5a.* Target gene expression was normalized to actin and made relative to WT uninjected expression levels. Injections were performed with 100pg of *wnt8* at the one cell stage. Expression of *axin2*, *nkd1*, and *sp5a* tended to increase in *axin2*^-/-^ mutants at both 30% and 50% epiboly but only *axin2* expression increased significantly at 30% epiboly. However, in the *nkd1*^-/-^ and *axin2*^-/-^;*nkd1*^-/-^ mutants, *axin2* and *nkd1* expression did not increase substantially compared to wild type. In contrast, *sp5a* expression showed a significant increase at 30% and 50% epiboly in the *nkd1*^-/-^ mutants and at 50% in the *axin2*^-/-^;*nkd1*^-/-^ (A-E, error bars represent SEM, * = p-value < 0.05, ** = p-value < 0.01, one-way ANOVA). Injection of Wnt8 into wild type (column 3) is often significant to uninjected wild type (column 1), but this is not shown for clarity.

### Wnt regulator mutants have differential effects on *gsc* expression

To explore this differential effect further, we chose to look at the expression of *gsc* by whole mount *in situ* hybridization (WMISH) to determine how the dorsal organizer is affected by loss of Wnt feedback regulation. The dorsal organizer is under the influence of both maternal and zygotic canonical Wnt signalling^61-62^. From dome to 50% epiboly, the arc of *gsc* expression on the dorsal side of the embryo can be quantified as a read out for Wnt activity. This would also determine if there were an expansion of Wnt signaling or ectopic Wnt signaling. At dome stage and 30% epiboly there was no significant difference in the arc of *gsc* expression in the *axin2*^-/-^ and *nkd1*^-/-^ maternal-zygotic mutants when compared to wild type (Fig. 5A,B). In contrast, at 50% epiboly, the *axin2*^-/-^ maternal-zygotic (TL background) embryos had an 8.1° increase in *gsc* arc angle which was not seen in either the *nkd1*^-/-^ maternal-zygotic (TU background) or *axin2*^-/-^;*nkd1*^-/-^ maternal-zygotic embryos (predominately TU background) (Fig. 5C,D).

**Figure 5.**
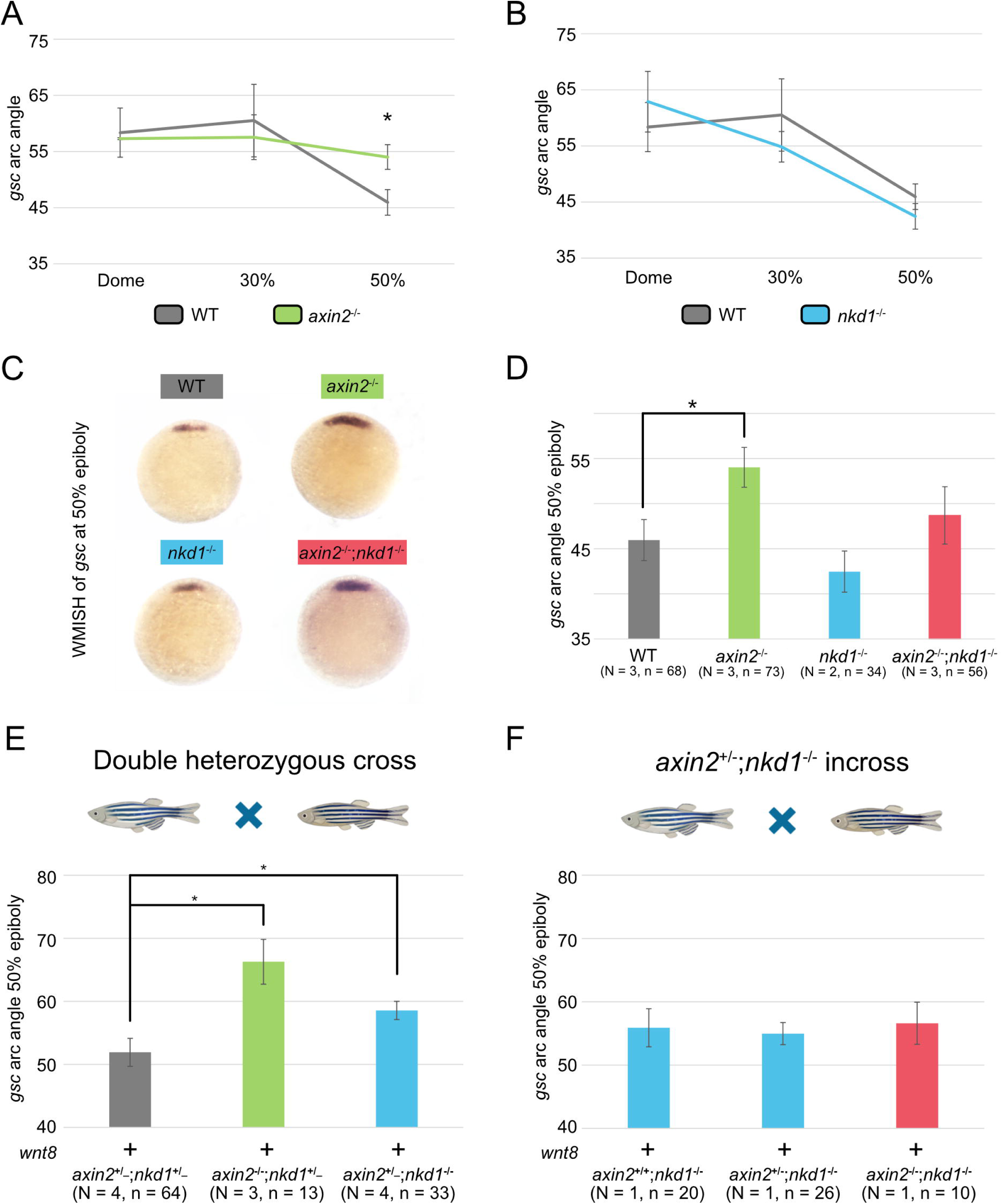
Wnt regulator mutants have differential effects on *gsc* expression. (A-D) WMISH for *gsc* was performed on embryos at 3 different stages for *axin2*^-/-^ and *nkd1*^-/-^ single mutants (A,B) as well as 50% epiboly for all Wnt regulator mutants (C) and measured for the arc of *gsc* expression (D). At 50% epiboly, the arc angle of *gsc* expression was significantly broader in the *axin2*^-/-^ embryos, which was not seen in either *nkd1*^-/-^ or *axin2*^-/-^;*nkd1*^-/-^ mutants. (E-F) Analysis of *gsc* expression on embryos at 50% epiboly injected with 100pg of *wnt8* from either an *axin2*^+/-^;*nkd1*^+/-^ incross (E) or an *axin2*^+/-^;*nkd1*^-/-^ incross (F). (E) The *axin2*^-/-^;*nkd1*^+/_^ and *axin2*^+/_^;*nkd1*^-/-^ embryos had an increase in *gsc* arc angle when compared to *axin2*^+/_^;*nkd1*^+/_^. (F) In contrast, *axin2*^-/-^;*nkd1*^-/-^ double mutant embryos showed no increase in *gsc* arc angle when compared to the single knockout genotypes. Injections were performed at the one cell stage and arc angle measurements were taken on ImageJ. Embryos were genotyped using indel detection by amplicon analysis and PCR based methods. A one-way ANOVA was used for all statistics with * = p-value < 0.05, error bars represent SEM (A,B,C,E) and standard deviation (F).

This was further validated by knocking down Axin2 using morpholinos which caused an increase in *gsc* arc angle at 50% epiboly but not at 30% epiboly (Supplemental fig. 7). This is in contrast to previous research from our lab where knocking down Nkd1 using a morpholino increased *gsc* expression at 30% epiboly^20^. This may be a consequence of knocking down translation via morpholino versus knocking out maternal and zygotic transcription in the *nkd1*^-/-^ or *axin2*^-/-^;*nkd1*^-/-^ knockouts shown here. This suggests that Axin2, but not Nkd1, normally functions to restrict the expression of *gsc*.

To better evaluate if the genetic background is a contributing factor (for example, the TL may be more sensitive than the TU background), we overexpressed *wnt8* in embryos that were generated from a double heterozygote incross (equal parts TU and TL), processed for WMISH, scored for phenotype, and then genotyped. Unlike the *axin2*^-/-^;*nkd1*^-/-^ maternal-zygotic mutants above, these embryos would be zygotic mutants for all 3 genotypes with an approximately equal contribution of the TL and TU genetic backgrounds. Embryos with Axin2 knocked out (*axin2*^-/-^;*nkd1*^+/_^) had a 14.4° increase in *gsc* arc angle, whereas, embryos with Nkd1 knocked out (*axin2*^+/_^;*nkd1*^-/-^) had a 6.6° increase in *gsc* arc angle when compared to embryos with at least one wild type allele of both Axin2 and Nkd1 (*axin2*^+/_^;*nkd1*^+/_^), both of which were significant (Fig. 5E). Similarly to the results in the *axin2*^-/-^ maternal-zygotic (TL background), the *axin2*^-/-^;*nkd1*^+/_^ (TU/TL background) had the greatest increase in *gsc* arc angle with *wnt8* overexpression and does not support the notion that the TL background is more sensitive.

The evaluation of *axin2*^-/-^;*nkd1*^-/-^ from the double heterozygote incross was hindered by the low numbers that were available, as only 1/16 of all offspring from a double heterozygous cross would be *axin2*^-/-^;*nkd1*^-/-^. Therefore, to increase the number of *axin2*^-/-^;*nkd1*^-/-^ embryos, *wnt8* overexpression was performed on embryos from a *axin2*^+/-^;*nkd1*^-/-^ (predominantly TU background) incross. Twenty five percent of these embryos would be zygotic *axin2*^-/-^ and maternal-zygotic *nkd1*^-/-^. Similar to the maternal-zygotic mutants experiments above (Fig. 5A-D), *wnt8* injected *axin2*^-/-^;*nkd1*^-/-^ mutants did not have an increase in *gsc* (Fig. 5F). Collectively, this suggests that loss of Nkd1 rescues the loss of Axin2 or that loss of Nkd1 is epistatic to loss of Axin2 and further suggests that the effect of loss of Axin2 on *gsc* expression appears to be independent of genetic background and maternal sources.

The evidence thus far suggests that there are at least 3 different responses in the loss of the negative feedback regulators: 1) Additive: terminal neuromasts are more severe in *axin2*^-/-^;*nkd1*^-/-^ mutant; 2) Counteractive: *axin2*^-/-^;*nkd1*^-/-^ rescues Wnt8 overexpression in single mutants and 3) Nkd1 epistasis: *sp5a* expression is up in *nkd1*^-/-^ and *axin2*^-/-^;*nkd1*^-/-^ mutants but not in *axin2*^-/-^; *gsc* and *axin2* expression up in *axin2*^-/-^, but there is no change in *nkd1*^-/-^ or *axin2*^-/-^;*nkd1*^-/-^. Further, we only observed the counteractive response in the eyeless phenotype, suggesting that this may be related to other signaling phenomenon occurring during patterning of the eye field.

### RNA-seq analysis suggests *axin2*^-/-^;*nkd1*^-/-^ mutants are more like *nkd1*^-/-^

To better understand the relationship between these two Wnt regulators, we performed RNA-sequence analysis on embryos at 50% epiboly, a time point when we observed significant effects on gene expression in the mutants. In addition to the overall analysis in gene expression, we also attempted to identify groups that reflect the three different responses identified above.

RNA-sequencing was performed on the Wnt regulator mutants as well as the two wild-type background genotypes: TU and TL. The two genetic backgrounds were useful in identifying and removing genetic variability unrelated to the mutations. Three biological replicates were isolated on different days, but from the same parents for each genotype. RNA-sequencing was performed using paired-end reads at a depth of 50 million reads.

A principal component analysis was performed on all genes. This revealed that all biological replicates were closely related to each other demonstrating the reproducibility of the dataset. Further, the *axin2*^-/-^ mutants were more similar to the TL background from which it was derived, while the *nkd1*^-/-^ mutants clustered closely with the *axin2*^-/-^;*nkd1*^-/-^ double mutants (Fig. 6A), which is consistent with the predominantly TU background in the double mutants. Furthermore, a heat map of the top 1000 differential regulated genes also shows a similar trend where *axin2*^-/-^;*nkd1*^-/-^ embryos are more similar to *nkd1*^-/-^ (Fig. 6B). This data fits very well with the *sp5a* expression we observed above. To further explore the data, Venn diagrams were created to identify overlap in differentially expressed genes (DEG) found in each of the Wnt regulator mutants. Before the Venn diagrams were created, genes that were differentially expressed (FDR < 0.05) between TU and TL were removed to strengthen the reliability of our data by removing genes that were variable between the background genotypes. Using an absolute log2FC >2 and FDR < 0.05 filtering method revealed 735 DEG in the *axin2*^-/-^ mutants, 869 DEG in *nkd1*^-/-^ mutants and 312 DEG in the *axin2*^-/-^;*nkd1*^-/-^ (Fig. 6C,D). We attributed the reduced number of DEG in *axin2*^-/-^;*nkd1*^-/-^ mutants to the fact that they were filtered against both the TU and the TL backgrounds, while the single mutants were only filtered against their respective backgrounds.

**Figure 6.**
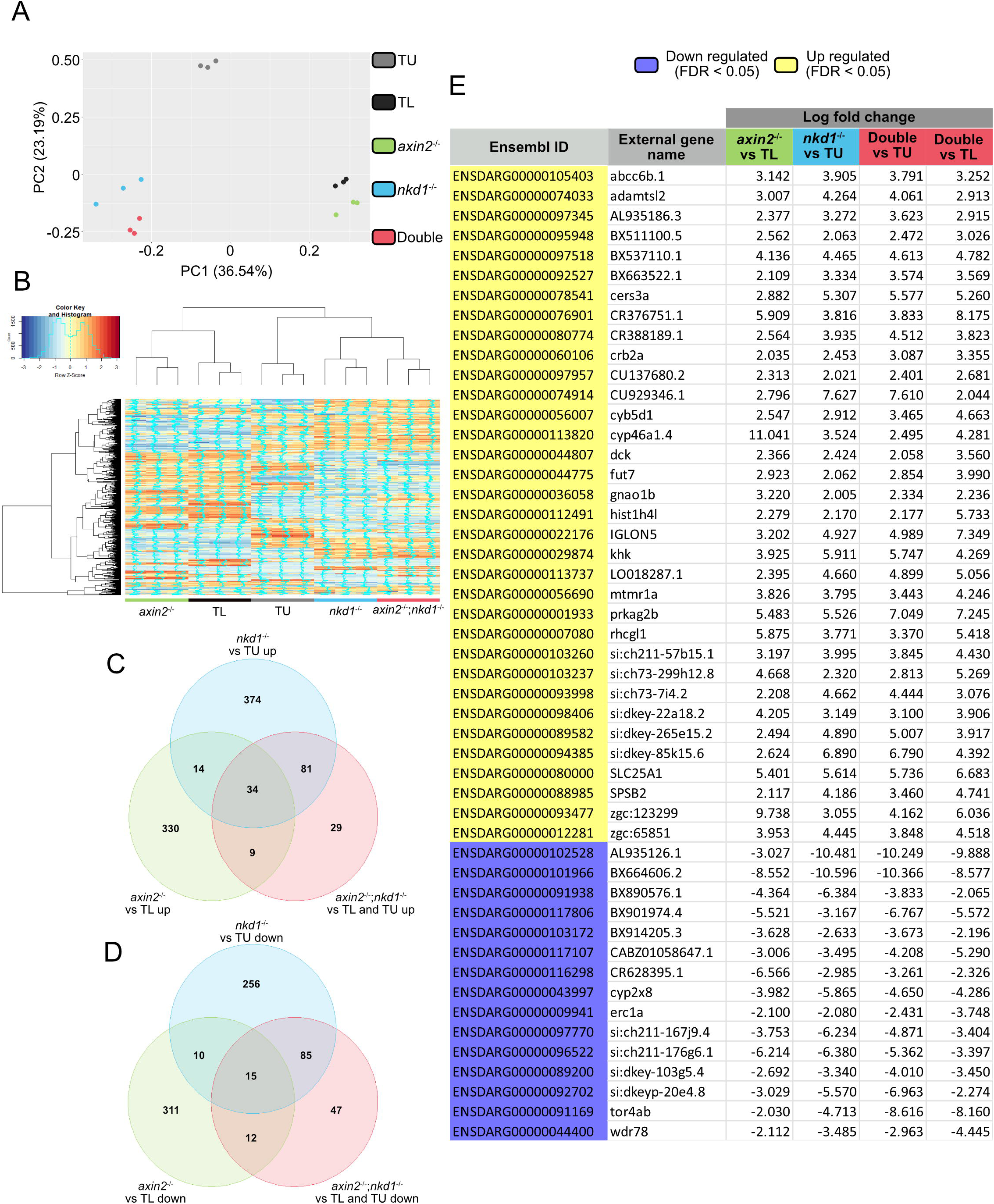
*axin2*^-/-^;*nkd1*^-/-^ embryos have a more similar RNA profile to *nkd1*^-/-^ than to *axin2*^-/-^. RNA-seq was performed at 50% epiboly for all Wnt regulator mutants as well as two wild type background genotypes: TU and TL. (A) PCA analysis and a (B) heat map analysis of the top 1000 differentially regulated genes shows that that *axin2*^-/-^;*nkd1*^-/-^ double mutant embryos have a similar RNA profile to the *nkd1*^-/-^ single mutant embryos. (C-D) Venn diagrams were made for upregulated genes (C) (log2FC > 2 and an FDR < 0.05) and down regulated genes (D) (log2FC < -2 and an FDR < 0.05) with genes that were significantly different (FDR < 0.05) between TL and the TU backgrounds being removed. (E) A gene list for the 34 genes that were upregulated and the 15 genes that were down regulated for each of the Wnt regulator mutants was made with the log fold change values provided for comparison. For more information on DEG for each Wnt regulator mutant refer to supplemental tables 6-9.

From this, we conclude that the clustering of the *nkd1*^-/-^ mutants with *axin2*^-/-^;*nkd1*^-/-^ and both of these away from the *axin2*^-/-^ mutants is not due to the number of genes that are affected. Instead, it may be due to gene expression levels found in the mutants. Although not significant, a comparison of the expression values of the 34 upregulated genes shared in all mutant genotypes demonstrated that loss of Nkd1 had a greater impact on gene expression levels compared to loss of Axin2 (Supplemental fig. 8). A similar trend was observed in the 15 genes they had in common that are downregulated. Importantly, this was only observed for the shared genes, as comparison of expression levels of all the DEG for each genotype revealed nearly identical violin plots (Supplemental fig. 8C).

While both single mutants each have several hundred DEGs we were still perplexed by the lack of overlap in the number of shared genes. We wanted to determine if this was a consequence of reduced expression levels of genes in the *axin2*^-/-^ that were not shared. That is, do the 735 genes (outside of the 34 shared ones) found to be differentially expressed in *axin2*^-/-^ have differential expression in the *nkd1*^-/-^ mutant but just don’t reach the threshold of significance? To test this, we evaluated the raw expression levels of the 735 DEG identified in the *axin2*^-/-^ mutants in the *nkd1*^-/-^ dataset. For comparison, we also performed the reciprocal analysis (Supplemental Fig. 9). This analysis demonstrates that the non-overlapping DEG in the *axin2*^-/-^ remain relatively unchanged in the *nkd1*^-/-^ and *axin2*^-/-^;*nkd1*^-/-^ double mutants (Supplemental fig. 9A,B). This suggests that modifications to the transduction of the Wnt signal in the *axin2*^-/-^ mutant are very different to the modifications that occur with the loss of Nkd1 and supports the notion that these modifications are epistatic to the *axin2*^-/-^ modifications.

Of the 34 common genes that were upregulated and 15 common genes that were downregulated in all the Wnt regulator mutant genotypes, a large portion (28) of the 49 differentially expressed genes were uncharacterized with the vast majority of these (11/15) being downregulated (Fig. 6E).

To determine if the RNA-seq data supports the “additive” response group, we identified genes that were up (or down) in both *axin2*^-/-^ and *nkd1*^-/-^ mutants by Log2FC > 1 whose expression was exacerbated up (or down) in *axin2*^-/-^;*nkd1*^-/-^ mutants. This identified one gene, *Prkag2b*, that had a 5.5 fold change in both *axin2*^-/-^ and *nkd1*^-/-^ mutants and a 7.0 and 7.2-fold increase in the *axin2*^-/-^;*nkd1*^-/-^ vs TU and vs TL, respectively. There were 8 genes (Supplemental table 1) that were additive in their reduced expression.

To determine if there was evidence for the “counteractive” response, we identified genes that were up (or down) by Log2FC > 1 in both the single mutants (same direction) but were either unchanged or down (or up) by less than 1-fold in the *axin2*^-/-^;*nkd1*^-/-^mutants. This identified seven genes (Supplemental table 2), all of which had unchanged expression in the *axin2*^-/-^;*nkd1*^-/-^ mutants.

To determine how many genes were in the “Nkd1 epistasis” group, we selected genes that were differentially expressed by an absolute Log2FC > 1 in both the *nkd1*^-/-^ and *axin2*^-/-^;*nkd1*^-/-^ mutants, but were unchanged in the *axin2*^-/-^ mutants. This identified 237 genes. Increasing the stringency to absolute Log2FC > 2 resulted in 74 genes and a further increase to absolute Log2FC > 3 resulted in 33 genes (Supplemental table 3). Thus, while there is no significant support for “additive” or “counteractive” responses in the RNA-seq data, there is clearly support for the Nkd1 epistasis response. We also determined if there was an “Axin2 epistasis” group that was differentially expressed in *axin2*^-/-^ and *axin2*^-/-^;*nkd1*^-/-^ by an absolute Log2FC > 1, but were unchanged in the *nkd1*^-/-^ mutant and identified only five genes (Supplemental table 4).

### Wnt target gene expression in *axin2*^-/-^;*nkd1*^-/-^ mutants is more similar to *nkd1*^-/-^ than *axin2*^-/-^

Analysis of the 237 “Nkd1 epistasis” genes by GO and Panther analysis revealed no significant hits, regardless of the stringency we used. Curiously, only 3 of the 237 genes were related to Wnt signaling: *gsk3b*, *fzd10* and *caveolin*. We performed similar GO analyses on each of the three mutant genotypes and in each case, there were no significant hits. Analysis of the non-significant hits did not reveal any patterns; however broad neuronal systems were abundant. Surprisingly, Wnt signaling did not appear in any of the GO analyses which could be due to the analysis focusing on the proteins in the pathway rather than the expression of Wnt target genes (data not shown).

Despite the high levels of *sp5a* identified in our qRT-PCR analysis, this gene was not identified as being a significant hit in the RNA-seq data. Thus, we wanted to investigate this gene and other known Wnt target genes. Using the RNA-seq data, a list was created of 59 known Wnt target genes identified by the Wnt signaling community. It should be noted that many of these are not unique to Wnt signaling and may have a more significant role in other pathways. Of the 59 Wnt target genes, 15 of them had differential expression in *axin2*^-/-^;*nkd1*^-/-^ mutant embryos (FDR < 0.05 vs TU and FDR < 0.05 vs TL). Of these 15 differentially expressed genes, nine of them had similar expression to *sp5a* seen earlier by qRT-PCR: *fsta, lef1, nkd1, six3b, sox17, sox2, sp5a, tbx3a, and tiam1a* (Fig. 7). The inclusion of *sp5a* in the list is validation of the qRT-PCR data. Furthermore, the levels of *axin2* and *nkd1* in the RNA-seq data are also consistent with our qRT-PCR analysis, showing modest but consistent increases in expression in all three mutant genotypes.

**Figure 7.**
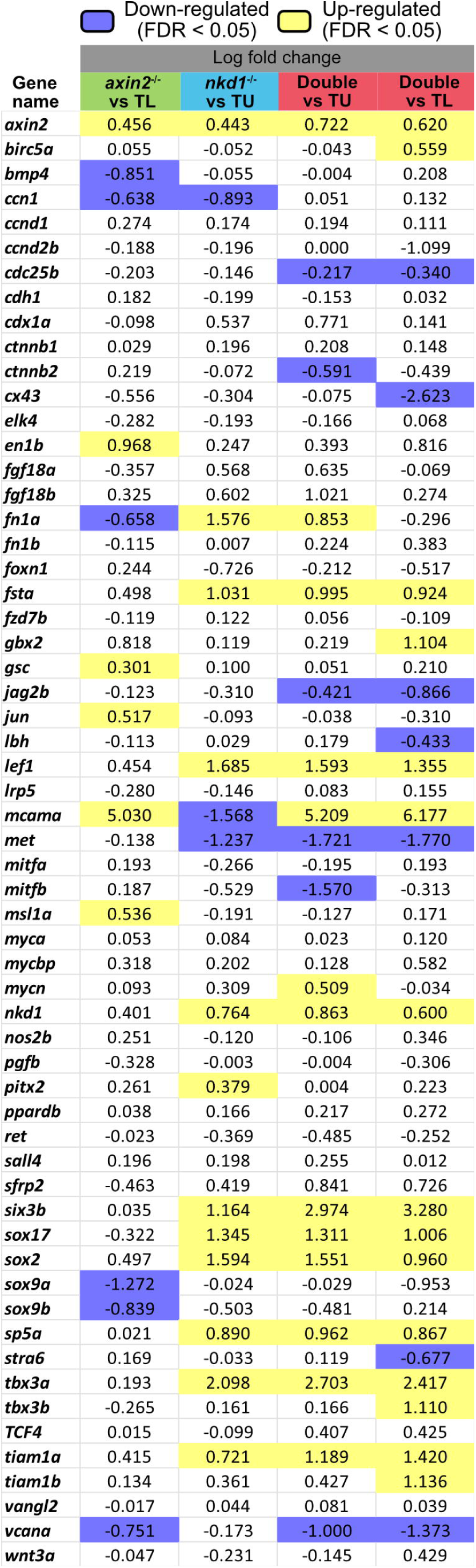
Wnt target gene expression in *axin2*^-/-^;*nkd1*^-/-^ is more similar to *nkd1*^-/-^ than *axin2*^-/-^. 59 Wnt target genes identified by the Wnt signaling community were used to create a list of expression comparisons between the Wnt regulator mutants compared to their respective backgrounds using the RNA-seq data.

Collectively, the differential gene expression analysis suggests that loss of Nkd1 has a greater impact on gene expression levels compared to the loss of Axin2 and that there is no additive, synergistic or counteractive effect in the loss of both Nkd1 and Axin2. While the GO analysis did not reveal any significant hits, there was a modest, but consistent effect on Wnt target genes. Further, it seems that there are a significant number of uncharacterized genes that are targets of the Wnt pathway that have yet to be described, highlighting the complexity and depth of this pathway in multipotent progenitor cells.

The lack of overlap in target genes between *axin2*^-/-^ and *nkd1*^-/-^ (only 73 genes out of a combined 1,531) is curious but does support our qRT-PCR and *gsc* expression analysis. This suggests that each contribute to modifications of β-catenin-mediated signaling resulting in an altered transcriptional response, which may include alternative transcriptional start sites in their shared genes. To explore this further, we evaluated differential transcript usage (DTU) in the different genetic backgrounds but found no obvious pattern for DTU when comparing the Wnt regulator mutants. Several of the most significant hits are shown in supplemental figure 10. Accordingly, we did not pursue this further.

### Knocking out Axin2, but not Nkd1, increases cytoplasmic levels of β-catenin

As both Axin2 and Nkd1 interact with β-catenin in the cytoplasm, we next wanted to determine if there were changes in the levels of cytoplasmic and nuclear β-catenin. As Axin2 is predicted to target β-catenin for ubiquitin mediated degradation, we predicted that loss of Axin2 would result in more cytoplasmic β-catenin. In contrast, we have previously demonstrated that Nkd1 functions to inhibit the nuclear accumulation of β-catenin and so its loss is predicted to result in more nuclear β-catenin but not necessarily a change in cytoplasmic levels^22^. To test this, we first performed Western blot analysis on cytoplasmic fractions from 30% epiboly embryos, a time point where we have previously shown cytoplasmic β-catenin levels to be sensitive to Wnt signaling hyperactivation^22^. Further, Pan-cadherin was used as a control to ensure that the cytoplasmic fractions were free of plasma membrane as β-catenin is localized to the membrane due to its interactions with cadherins^63^. Compared to uninjected controls, wild type embryos had a relative cytoplasmic β-catenin increase of 1.86 when overexpressed with *wnt8*, which is consistent with our previous reports (Fig. 8A)^22^. Analysis of β-catenin levels in the Wnt regulator mutants demonstrated a significant increase in *axin2*^-/-^ mutants. In particular, *axin2*^-/-^ and *axin2*^-/-^;*nkd1*^-/-^ had a relative cytoplasmic β-catenin increase of 2.90 and 3.52, respectively, compared to their uninjected genotype control (Fig 8B). These were significantly higher than the 1.86 increase seen in wild type *wnt8* injected embryos. By comparison, the *nkd1*^-/-^ mutant had a relative cytoplasmic β-catenin increase of 2.17, which was not significant when compared to wild type (Fig. 8B) but supports the higher levels seen in the *axin2*^-/-^;*nkd1*^-/-^ mutant. This supports the model that Axin2 affects cytoplasmic levels of β-catenin^17,22,31^. While the statistics support our model that loss of Nkd1 does not alter cytoplasmic β-catenin levels, we are not convinced that there is no change in *nkd1*^-/-^ mutants, especially when *axin2*^-/-^;*nkd1*^-/-^ shows a modest additive effect.

**Figure 8.**
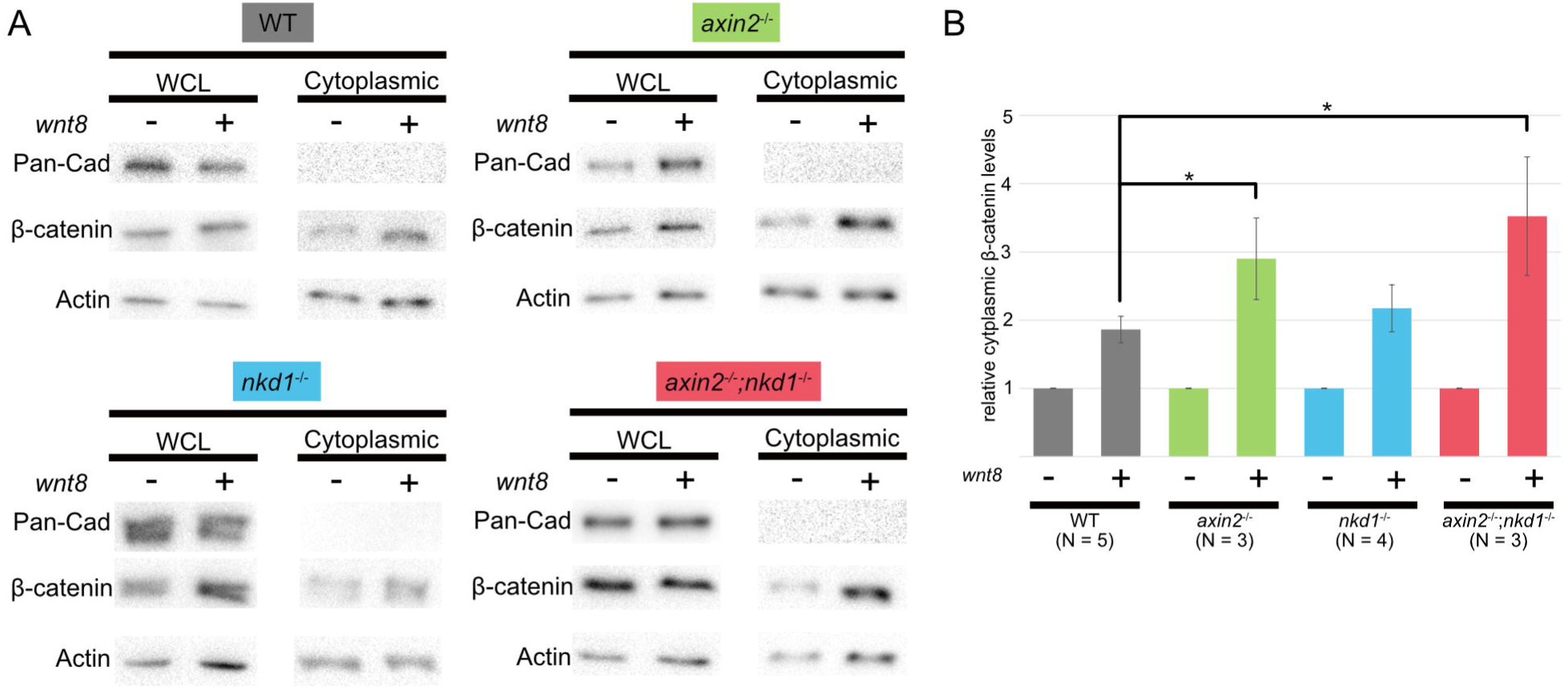
Knocking out Axin2, but not Nkd1, increases cytoplasmic levels of β-catenin. (A) Whole cell lysate and cytoplasmic fractions of embryos from wild type and Wnt regulator mutants, with or without 200pg of *wnt8* injected, were collected at 30% epiboly for Western blot analysis for cytoplasmic levels of β-catenin. Pan-cadherin was used to confirm purity of the cytoplasmic fraction and Actin was used as a loading control. (B) Since each genotype was run on separate western blots, the increase in cytoplasmic β-catenin when *wnt8* was overexpressed was made relative to their uninjected counterpart. Both *axin2*^-/-^ and *axin2*^-/-^;*nkd1*^-/-^ embryos had a greater increase in cytoplasmic β-catenin levels with *wnt8* overexpression when compared to wild type embryos. Error bars represent SEM, * = p-value < 0.05, one-way ANOVA.

### Wnt regulator mutants have no change in nuclear β-catenin levels at 30% epiboly

With the increase in cytoplasmic β-catenin in at least the *axin2*^-/-^ mutants, we next wanted to evaluate the levels of β-catenin in the nucleus. In our experience, nuclear fractions cannot be sufficiently purified from β-catenin-containing membrane fractions in zebrafish embryos for Western blot analysis, therefore we chose to perform immunohistochemistry at 30% epiboly using anti-β-catenin antibodies^20,22^. We first evaluated nuclear β-catenin levels in the different genetic backgrounds along the ventrolateral domain which has endogenous Wnt signaling and nuclear β-catenin. In contrast, cells outside of the ventrolateral domain were assessed for potential ectopic nuclear β-catenin. While difficult to quantify levels between embryos, we did not observe any obvious differences in nuclear β-catenin levels when compared to wild type embryos (Supplemental fig. 11).

### Knocking down Axin2 and/or Nkd1 does not increase nuclear β-catenin levels when Wnt signaling is hyperactive

We next set out to evaluate nuclear β-catenin in the presence of exogenous Wnt8. To do this, we injected a cocktail of Axin2 and/or Nkd1 sgRNAs, Cas9, and *wnt8* along with a tracer molecule (Dextran or *mcherry*) into 1 cell of an 8-cell stage wild type blastocyst to generate mosaic crispants. Embryos were fixed at 30% epiboly and immunohistochemistry was performed with anti-β-catenin antibodies. Images were captured away from the ventrolateral domain to avoid any influence from endogenous Wnt signaling. Using this design, labelled cells would be Wnt8 and crispant positive, but juxtaposed cells would be crispant negative, but still receive the Wnt ligand^22^. Thus, we compared nuclear β-catenin levels in label-positive cells with juxtaposed label-negative cells. Knocking down Axin2 and/or Nkd1 did not have any significant difference in nuclear β-catenin levels when compared to uninjected juxtaposed cells (Fig. 9). This is in contrast to previous findings where knocking down Nkd1 in the dorsal forerunner cells of zebrafish, and knocking out Axin2 in mice, caused an increase in nuclear β-catenin^33,64^. It is possible that our assay is not sensitive enough to detect subtle differences considering we are knocking down regulators of the pathway and not essential components. It is also possible that the sgRNAs were ineffective, but the same sgRNA were used in the Wnt reporter analysis with success as well as their efficiency being validated by indel detection by amplicon analysis (not shown). Taken together, loss of Wnt regulation via Axin2 and Nkd1 results in an increase in cytoplasmic β-catenin (at least significantly for Axin2), but this does not translate into a measurable increase in nuclear β-catenin.

**Figure 9.**
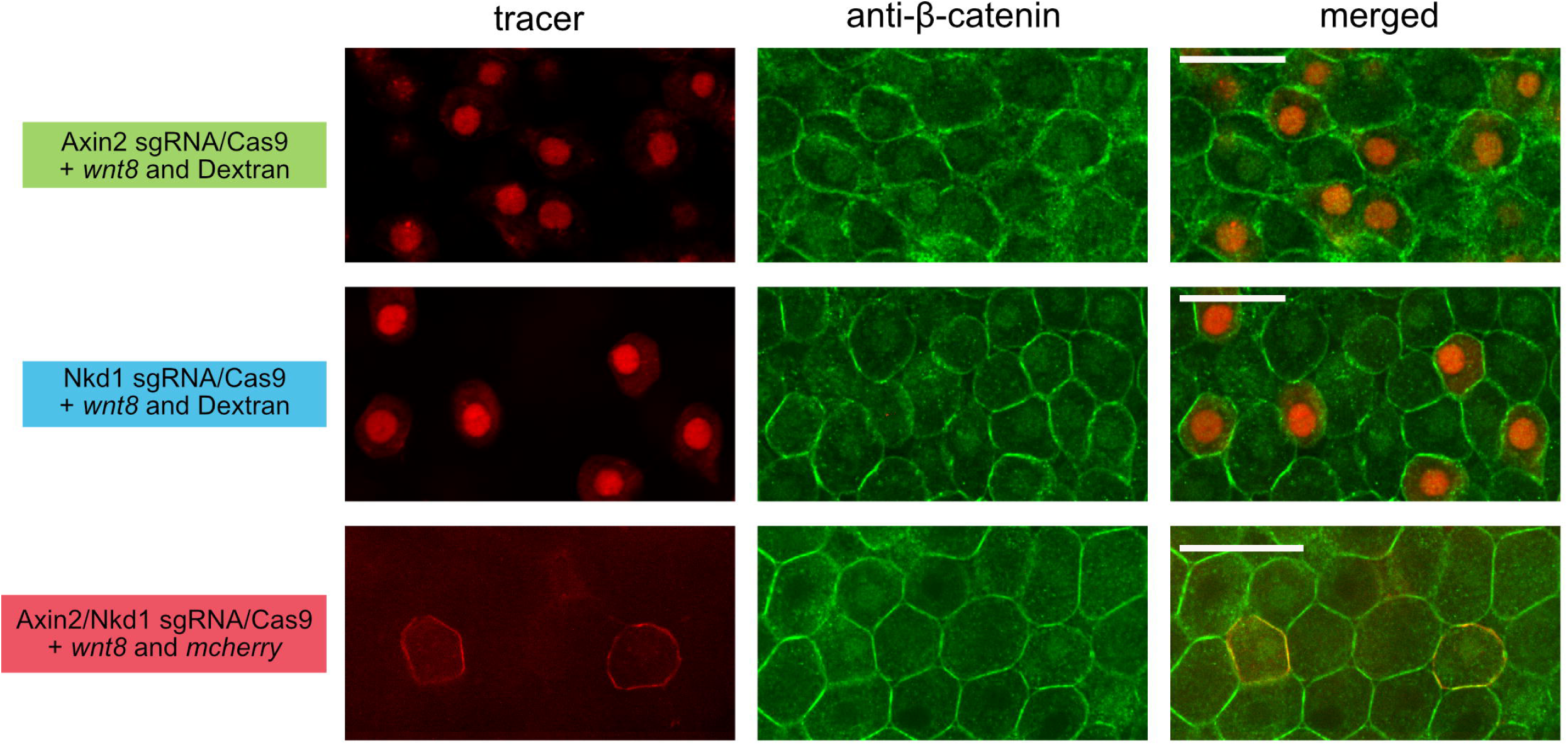
Knocking down Axin2 and/or Nkd1 does not increase nuclear β-catenin levels when Wnt signaling is hyperactive. Embryos were injected with a cocktail that contained Axin2 and/or Nkd1 sgRNA/Cas9, *wnt8,* and a tracer molecule (Dextran or mcherry) into 1 cell of an 8-cell stage embryo. Embryos were fixed at 30% epiboly and immunohistochemistry was performed for β-catenin. When Axin2 and/or Nkd1 was knocked down, there was no increase in nuclear β-catenin intensity when compared to juxtaposed uninjected cells. Scale bar = 20µM.

### Wnt regulator mutants have lower expression of metabolism proteins

The differences that we have observed in phenotypes, gene expression and cytoplasmic β-catenin levels suggests that Axin2 and Nkd1 have both unique and overlapping ways of controlling Wnt signaling and that this might be reflected in the protein complement in each of the mutants. We first tried to purify β-catenin to identify proteins that might be differentially interacting with it under the different genetic conditions; however, we were unable to purify sufficient levels of β-catenin from zebrafish embryos to perform this analysis. Instead, we decided to evaluate the entire protein pool using mass spectrometry with 30% blastocysts that have been rigorously purified away from yolk proteins. Using this strategy, we identified differentially expressed proteins between the Wnt regulator mutants. These results were variable between biological replicates which, despite our best efforts, was likely due to yolk contamination during isolation. Therefore, we chose samples with similar ionic chromatograms to perform our analysis (Fig. 10A). From this, we discovered 53 differentially expressed proteins (FDR < 0.1, logFC ≥ 1, peptides ≥ 2) in the Wnt regulator mutants with the majority of the proteins being either ribosomal subunits or related to metabolism (Fig. 10B). Furthermore, and similar to the RNA-seq data, we see that the *axin2*^-/-^;*nkd1*^-/-^ mutant is more similar to the *nkd1*^-/-^ mutant, whereas the *axin2*^-/-^ mutant is similar to wild type; however, the proteins that were differentially expressed in mass spectrometry did not always correlate to differential expression in the RNA-seq results. Furthermore, in the *nkd1*^-/-^ and *axin2*^-/-^;*nkd1*^-/-^ mutants, there is an overall trend of decreased protein expression levels. Of the 53 proteins, there were 18 proteins (Fabp3, Aldh2.2, Prdx1, ldhba, Pcxa, Tcp1, Igf2bp3, Pkma, Actb2, Cct5, Prdx6, Tpi1a, Prdx2, Ppa1b, Sod1, Cox17, Ranbp1, Stmn1a) that have a connection to Wnt signaling (Fig. 10C). Interestingly, most of the proteins related to Wnt have been demonstrated to stimulate Wnt signaling, as knocking out/down the proteins leads to either reduced β-catenin levels or Wnt target gene expression. Finally, using string analysis, we identified the top KEGG pathways to be ribosomal proteins and pyruvate metabolism related proteins (Fig. 10D-E). While this technique requires refinement to generate high quality data it does support our model where Axin2 and Nkd1 have both overlapping and unique effects in regulating Wnt signaling.

**Figure 10.**
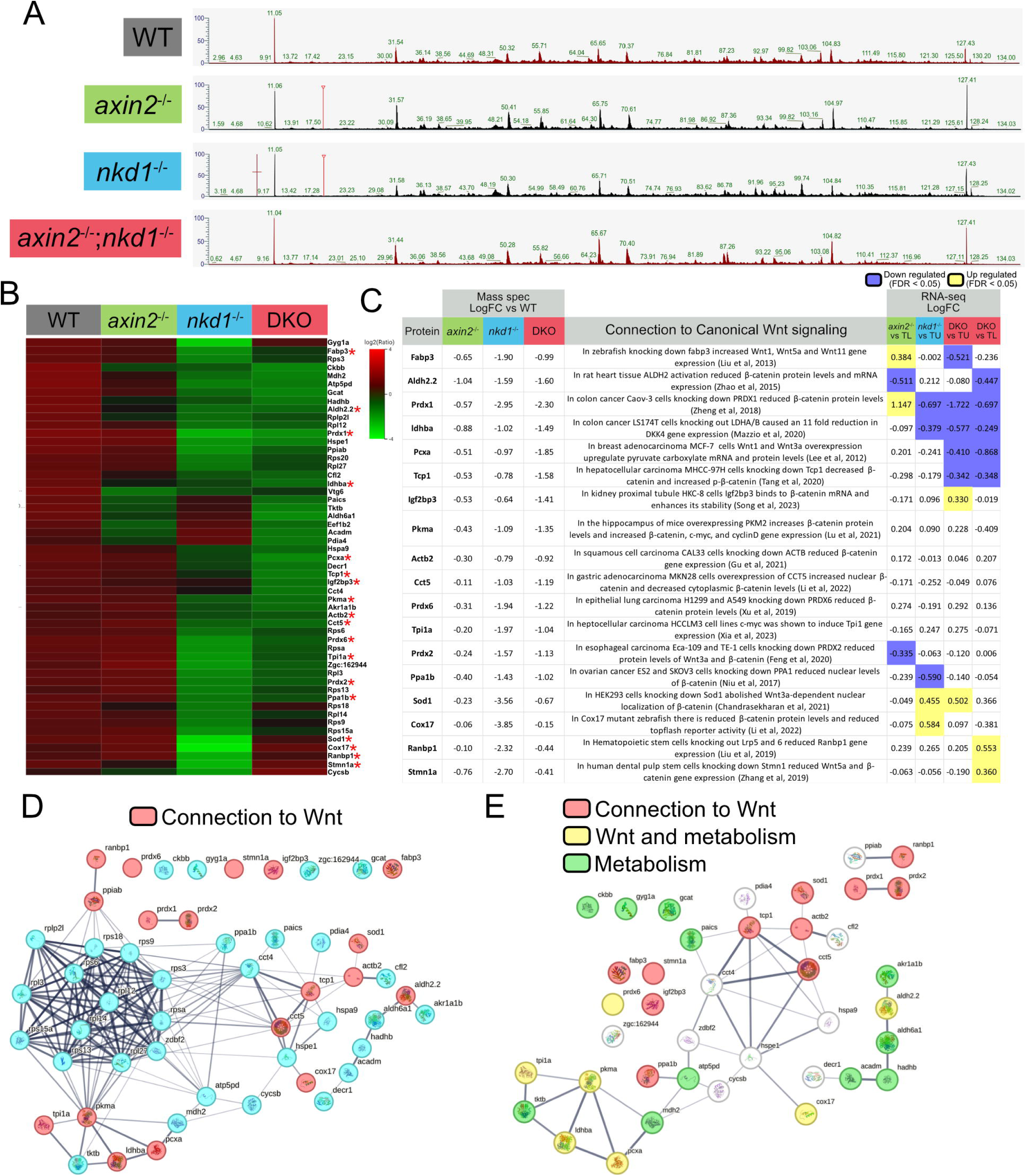
Wnt signaling mutants have lower expression of metabolism proteins. Mass spectrometry was performed on embryos at 30% epiboly. (A) Only samples with similar total ionic chromatograms were used, resulting in only 1 biological replicate for each genotype. (B) Heat map of 53 differentially expressed proteins using PEAKS 10 software (FDR < 0.1, logFC ≥ 1, peptides ≥ 2). (C) Eighteen proteins were identified to have a connection to Wnt signaling in the literature: Fabp3^87^, Aldh2.2^88^, Prdx1^75^, Idhba^89^, Pcxa^90^, Tcp1^91^, Igf2bp3^92^, Pkma^93^, Actb2^94^, Cct5^95^, Prdx6^96^, Tpi1a^97^, Prdx2^76^, Ppa1b^98^, Sod1^99^, Cox17^100^, Ranbp1^101^, Stmn1a^102^. (D-E) String protein networks were created using all proteins identified (D) and a network of the 39 non ribosomal proteins (E).

## Discussion

Our initial hypothesis was that Axin2 and Nkd1 uniquely and independently affect Wnt/β-catenin signaling with Axin2 regulating the stability of cytoplasmic β-catenin, while Nkd1 prevents its nuclear accumulation^17,22^. We further anticipated a lack of a significant phenotype in each of the single mutants, based on the phenotypes in their mammalian counterparts^31,32,40,42^. Nonetheless, we speculated that with the different modes of action, there would be a convergence on the transcriptional targets and we anticipated a synergistic effect in the *axin2*^-/-^;*nkd1*^-/-^ double mutants resulting in more severe phenotype. Surprisingly, the phenotypic data was underwhelming. The heart looping data showed no evidence of epistasis or synergy. The axial and operculum defect in the *axin2*^-/-^ single and *axin2*^-/-^;*nkd1*^-/-^ double mutants suggests that Axin2 functions downstream of Nkd1, however, a detailed analysis clearly demonstrates that the *axin2*^-/-^;*nkd1*^-/-^ mutants resemble the *ndk1*^-/-^ mutants. This strongly suggests that Nkd1 functions downstream of Axin2 and that loss of Nkd1 is epistatic to loss of Axin2.

Evaluation of the entire transcriptome in a whole organism is a double edge sword. On one hand, we are evaluating the loss of genes in their native environment, which is very powerful. On the other hand, not every cell is actively engaged in Wnt signaling which can generate significant background. To maximize the former and minimize the later, we chose to evaluate the transcriptional readout of the mutants at 50% epiboly as there is substantial zygotic Wnt signaling occurring throughout the ventrolateral domain of the blastula. The transcriptional data supports Axin2 having a separate mode of Wnt antagonism from Nkd1. However, we don’t have any evidence that loss of Axin2 is epistatic to loss of Nkd1 or that Axin2 functions downstream of Nkd1. In contrast, the majority of our analysis supports the model that Nkd1 functions downstream of Axin2 and its loss is epistatic to the loss of Axin2 (Table 1).

**Table 1.** Summary of results. Each experiment is listed with the effects shown for each genotype. A (-) symbol signifies no observed effect, while the (+) symbol signifies an effect. **+** = somewhat affected, **++** = affected, **+++** = very affected. Colour coding reflects genotype. In the majority of our analysis the *axin2*^-/-^;*nkd1*^-/-^ mutant looks similar to *nkd1*^-/-^ but rarely has a more severe phenotype when compared to the single mutants.

| Experiment | *axin2*<sup>-/-</sup> | *nkd1*<sup>-/-</sup> | *axin2*<sup>-/-</sup>;*nkd1*<sup>-/-</sup> | *axin2*<sup>-/-</sup>;*nkd1*<sup>-/-</sup> is like? |
| --- | --- | --- | --- | --- |
| Heart looping | ++ | ++ | ++ | Both |
| Neuromast migration | ++ | ++ | +++ | Additive |
| Spine and operculum | ++ | - | ++ | *axin2*<sup>-/-</sup> |
| Thigmotaxis behavior | ++ | ++ | ++ | Both |
| Tap response | + | +++ | +++ | *nkd1*<sup>-/-</sup> |
| Eyeless Wnt8 | ++ | ++ | - | WT |
| Twisted axis Wnt8 | - | ++ | - | *axin2*<sup>-/-</sup> |
| Wnt activity in forebrain | ++ | + | ++ | *nkd1*<sup>-/-</sup> |
| Wnt activity in forebrain Wnt8 | +++ | + | ++ | *nkd1*<sup>-/-</sup> |
| BIO sensitivity | ++ | ++ | ++ | Both |
| *axin2* expression | ++ | + | + | *nkd1*<sup>-/-</sup> |
| *axin2* expression Wnt8 | ++ | + | + | *nkd1*<sup>-/-</sup> |
| *nkd1* expression | + | + | + | *nkd1*<sup>-/-</sup> |
| *nkd1* expression Wnt8 | + | + | + | Both |
| *sp5a* expression | - | +++ | +++ | *nkd1*<sup>-/-</sup> |
| *sp5a* expression Wnt8 | + | +++ | +++ | *nkd1*<sup>-/-</sup> |
| *gsc* expression at 50% | ++ | - | - | *nkd1*<sup>-/-</sup> |
| *gsc* expression with Wnt8 OE | ++ | ++ | ++ | Both |
| RNA-seq | + | ++ | ++ | *nkd1*<sup>-/-</sup> |
| RNA-seq Wnt target genes | + | ++ | ++ | *nkd1*<sup>-/-</sup> |
| Cytoplasmic β-catenin | ++ | - | ++ | *axin2*<sup>-/-</sup> |
| Nuclear β-catenin | - | - | - | WT |
| Nuclear β-catenin Wnt8 | - | - | - | WT |
| Protein expression | + | +++ | +++ | *nkd1*<sup>-/-</sup> |

The difficulty in working with negative feedback regulators is that they function to regulate the intensity, spatial distribution and/or duration of the signal and can act as a safety mechanism more than an integral component of the pathway, much like an airbag in a car^65^. For example, we see a more robust response in *sp5a* expression when we ectopically express Wnt8 in the mutants compared to wildtype. Nonetheless, we still observed significant increases in target gene transcription without the addition of exogenous Wnt8. We also speculate that the high number of DEG’s identified is an overestimation of the actual targets. Increasing the absolute Log2FC to >3 resulted in 33 genes that are significantly differentially transcribed in both the *nkd1*^-/-^ single and *axin2*^-/-^;*nkd1*^-/-^ double mutants but remained unchanged in the *axin2*^-/-^ mutant. One, Gsk3βb, is linked to Wnt signaling and is upregulated over 60-fold; two are related to transcriptional activity; one is a histone deacetylase and nine are uncharacterized proteins with the remainder having various unrelated functions. Similarly for *axin2*^-/-^ mutants with an absolute Log2FC>1 but remained unchanged in the *nkd1*^-/-^ or *axin2*^-/-^;*nkd1*^-/-^ mutants resulted in 52 genes, 1 of which is related to Wnt signaling (*sec12I8*); 9 are related to DNA binding; 4 are related to histone function; 10 are uncharacterized proteins and the remainder have various unrelated functions (Supplemental Table 5). The significant hits in DNA binding and histone modifications are in line with the model that loss of Axin2 and/or Nkd1 would alter Wnt signaling transcriptional activity, but this requires further investigation.

While the data is consistent with the loss of Nkd1 being epistatic to loss of Axin2, there are a few scenarios where we observe unexpected phenotypes in the presence of ectopic Wnt8. Firstly, Wnt8, but not BIO, induces a kinked tail phenotype in the *nkd1*^-/-^ mutants, which is rescued in *axin2*^-/-^;*nkd1*^-/-^ mutants. We speculate that the kinked tail phenotype is a result of perturbed Wnt/planar cell polarity (PCP) signaling, but how this is rescued by loss of Axin2 is unknown.

Secondly, Axin2 and Nkd1 are both expressed along the ventrolateral margin of the gastrulating embryo when the hindbrain is being patterned^20,54,66^. Predictably, the single mutants were highly sensitive to exogenous Wnt8, resulting in the classic Wnt gain of function eyeless phenotype. Our prediction was that the *axin2*^-/-^;*nkd1*^-/-^ double mutant embryos would be even more sensitive, but in fact the opposite was true; the *axin2*^-/-^;*nkd1*^-/-^ mutant was insensitive to exogenous Wnt8, resulting in a wild-type eye phenotype. To explain this, we must consider the role of Nkd1 in antagonizing the non-canonical Wnt/PCP pathway. We have previously demonstrated that Nkd1 functions to inhibit non-canonical Wnt/PCP signaling^20^. Furthermore, evidence suggests that there is cross talk between canonical Wnt and Wnt/PCP signaling where activation of Wnt/PCP can inhibit canonical Wnt/β-catenin signaling, however, this mechanism is not fully understood^67–70^. In this scenario, the presence of excess Wnt8 in combination with the loss of Nkd1 antagonism of Wnt/PCP during early gastrulation may lead to increased Wnt/PCP signaling and thus a higher inhibition of canonical Wnt signaling that could overcome the loss of Axin2 resulting in a rescue of the eyeless phenotype. In support of this model is the Wnt/PCP phenotype (kinked axis) that is induced only when Wnt8, and not BIO, is overexpressed in the *nkd1*^-/-^ mutants. Rescue of the eyeless phenotype through the loss of both Axin2 and Nkd1 suggests that the Wnt/PCP-Wnt/β-catenin interaction is occurring within the cytoplasm upstream of the destruction complex and is dependent on a Wnt ligand interacting with its receptor. As Dvls are involved in both Wnt/β-catenin and Wnt/PCP signaling^71^, bind Nkd1 and Axin1 but not Axin2^27^, act upstream of the destruction complex and are activated by Wnt ligands^72^, Dvls seem to be likely candidates to participate in this phenomenon but this requires further investigation.

Consistent with our results, there have been several studies investigating gene expression under different Wnt signaling scenarios, and Sp5 expression appears to be a highly upregulated Wnt target gene in these studies^55–57,60^. In addition, *sp5* expression is significantly higher than *axin2* and *nkd1* in these studies, which is also consistent with our study^55–57,60^. Similarly, when we compared our hits to the Wnt signaling study from Moya *et al*, 2014, we both identified *cyb5d1*, *wdr78*, *gnao1b*, and *crb2a* as Wnt target genes^56^. Further, we saw both upregulation and downregulation of Wnt target genes which is consistent with the research by Ewing *et al*, 2020 where Wnt target genes were also upregulated and downregulated when comparing HCT116 Δ45-β-catenin to HCT116 wild type cells^73^. This is also consistent with the function of β-catenin-dependent transcription being context dependent which is covered in depth in the review by Valenta *et al*, 2012^74^.

Our evaluation of the proteome by mass spectrometry is a unique approach as these studies are rarely performed on whole organisms, especially in zebrafish due to the high yolk protein content. While some replicates provided more hits, the poor reproducibility between replicates made it insufficient for any meaningful analysis and thus, we decided to focus on the samples with lower abundant proteins but with the most similar ion chromatogram profiles. This method identified 53 differentially expressed proteins with 18 of those proteins having known connections to Wnt signaling. For example, Prdx1 and Prdx2, proteins have been previously linked to enhancing β-catenin stability and were found to be downregulated in *nkd1*^-/-^ and *axin2*^-/-^;*nkd1*^-/-^ mutants in our results^73,75,76^. The transcript levels of *prdx1* were also significantly reduced, while *prdx2* transcripts were only slightly reduced. However, the majority of the proteins we identified were related to ribosomal assembly or metabolism, which may reflect the high translation rates occurring at 50% epiboly, a time when the embryo has fully transitioned into zygotic expression^77^. Interestingly, most of the proteins connected to Wnt signaling seem to have a promoting role in the pathway and are down regulated in the Wnt regulator mutants. This could suggest that the embryos are modulating metabolism to downregulate Wnt signaling when Axin2 and/or Nkd1 is/are knocked out. Thus, our results are consistent with Wnt signaling having a known role in metabolism at both the gene and protein levels, however this is far from comprehensive and will need to be investigated further^78^.

The differences between loss of Nkd1 and loss of Axin2 might suggest that Axin2 and/or Nkd1 are not negative feedback regulators of Wnt, but the overlapping results strongly suggests that they are which is consistent with the literature^17,22^. In particular, their sensitivity to Wnt8 overexpression or BIO treatment are well established indicators of excess Wnt signaling^19,21,79^. Furthermore, their common phenotypes in heart looping and neuromast migration and their effect on many Wnt target genes are also strong indicators that Axin2 and Nkd1 are involved in Wnt signaling.

Alternatively, it might be argued that the mutations in *axin2*^-/-^ and *nkd1*^-/-^ are not complete nulls, which would require reading through of the premature stop codons^80^. In support of this, we do not observe nonsense mediated RNA decay in the qRT-PCR data. Without specific antibodies to Axin2 and Nkd1, it is difficult to confirm a complete null, but there is evidence to suggest they are both nulls. First, phenotypes that are common to both have similar effects; for example, we observe similar effects in heart looping and sensitivity to exogenous Wnt8 for both mutants. We also observe a significant increase in *axin2* and *sp5a* expression in *axin2*^-/-^ and *nkd1*^-/-^ mutants, respectively, with and without exogenous Wnt8. These results were also validated using RNA-seq where we saw differential expression of Wnt target genes in the Wnt regulator mutants. Furthermore, the *axin2*^-/-^ mutant has a similar number of DEG compared to *nkd1*^-/-^. Finally, we have previously generated *axin2* and *nkd1* crispants using at least two unique, non-overlapping sgRNAs and evaluated their phenotypes and target gene expression with or without Wnt8. All Crispants had similar eyeless phenotypes and *axin2* and *nkd1* expression patterns as observed here^81^. Thus, while we cannot be 100% certain that we have complete nulls, we are confident that at the very least the majority of the Axin2 and Nkd1 proteins are non-functional. We also injected Nkd2 sgRNA/Cas9 into the *nkd1*^-/-^ mutants and found no evidence of compensation by Nkd2 (data not shown).

In conclusion, we have provided evidence that Nkd1 functions downstream of Axin2 and the destruction complex, as loss of Axin2 has minimal effect on Wnt signaling when Nkd1 is knocked out. Further, the evidence clearly demonstrates that Axin2 and Nkd1 uniquely modify Wnt signaling which we speculate happens at the level of β-catenin. How these modifications affect gene transcription will be an active area of investigation and provide important insights into the complex regulation of a signaling pathway involved in many development processes and diseases.

## Materials and methods

### Axin2 and Nkd1 Gene deletions

The website chop chop (https://chopchop.cbu.uib.no/) was used to design sgRNAs for Axin2 and Nkd1, with sgRNAs being selected based on stringency and location^82^. sgRNAs were made using the GeneArt Precision gRNA Synthesis Kit following the protocol provided in the kit (Invitrogen A29377, Supplemental table 10). The sgRNA were mixed with InvitrogenTM TrueCut Cas9 protein (Thermofisher) and injected at the one-cell stage. Cas9 efficiency was determined by indel detection by amplicon analysis.

### Neuromast staining

Embryos were treated with 0.003% PTU from 1dpf – 5dpf to inhibit pigmentation. At 5dpf larvae were fixed in 4% paraformaldehyde overnight at 4°C and then washed three times with PBST. Embryos were washed three times in alkaline tris buffer (0.5M Tris pH 9.5, 50mM MgCl2, 0.1M NaCl, 0.1% tween) and stained in alkaline tris buffer with 550nM Nitro blue tetrazolium chloride (NBT, Roche), and 400nM 5-Bromo-4-chloro-3-indolyl phosphate p-toluidine salt (BCIP, Roche).

### Microinjections

Injections were performed using a PV820 pneumatic PicoPump microinjector (World Precision Instruments) with needles pulled using a Model P-1000 Flaming Brown Micropipette Puller (World Precision Instruments, TW100F-4). Needles were cut to yield an injection volume of ∼1nL.

### BIO

Embryos were arranged into a 6 well dish at 4 hpf containing either 0µM, 0.25µM, 0.5µM, 1µM, and 1.5µM of BIO with 1% DMSO. Embryos were incubated at 28.5°C until 30 hpf and were fixed using 4% PFA overnight at 4°C. Images were taken and measurements for the area of the eye were done using ImageJ. The Wnt regulator mutants were treated in parallel with WT embryos. For the BIO treatment on Axin2 mixed embryos, an incross of *axin2*^+/-^ zebrafish was performed and embryos were treated with 0.75µM of BIO with 1% DMSO. Phenotypes for the eyes were recorded at 1dpf and each embryo was genotyped using fragment analysis.

### qRT-PCR

RNA was isolated from a pool of 10 embryos using The GENEzol™ TriRNA Pure Kit (FroggaBio). RNA was then directly used in the Luna® Universal One-Step RT-qPCR Kit (New England BioLabs). Primers listed in supplemental table 10.

### WMISH

Embryos were fixed at dome, 30%, and 50% epiboly using 4% paraformaldehyde overnight at 4°C. Embryos were washed three times in PBST and WMISH was carried out following the protocol provided in the Thisse and Thisse manuscript^83^.

### RNA sequencing preparation and analysis

RNA was isolated from a pool of 10 embryos at 50% epiboly using the GENEzol™ TriRNA Pure Kit (FroggaBio). RNA samples were DNase treated using the Invitrogen™ DNA-free™ DNA Removal Kit (Thermo Fisher Scientific) and an RNA integrity number of more than 8.0 was confirmed for all samples using the 4200 Tapestation system (Agilent). Poly(A) mRNA library preparation was performed using NEBNext Ultra II DNA library prep kit for illumina (New England BioLabs) and 2x100bp paired-end sequencing was performed at a depth of 50 million reads on an Illumina Novaseq 6000 platform by the University of Toronto Donelly Sequencing Centre. The reads were aligned using Rsubread v2.2.6^84^ to the Ensembl Genome Browser assembly ID: GRCz11. EdgeR v3.30.3^85^ was used to filter reads and DESeq2 v1.28.1^86^ was used to perform differential expression analysis.

### Western blotting

At 30% epiboly 10 embryos were manually deyolked using forceps and lysed using a 27-gauge needle in a TKM buffer (50mM Tris-HCl pH 7.5, 25mM KCl, 5mM MgCl2, 1mM EGTA, 0.02% sodium azide, 1X SIGMAFAST protease inhibitor cocktail tablets EDTA free). An aliquot was taken for whole cell lysate and the rest of the sample was centrifuged at 100,000 x G for one hour at 4°C. The supernatant containing the cytoplasm was collected, and the pelleted plasma membrane proteins was resuspended in TKM buffer. Westerns were performed using 1:1500 anti-β-catenin (BD Biosciences Cat# 610154), 1:2000 anti-Pan-cadherin (Abcam Cat# ab16505), and 1:5000 anti-Actin (Sigma Cat# A5441)

### Immunohistochemistry

At 30% epiboly embryos were fixed in 4% paraformaldehyde at 4°C overnight. Embryos were washed three times in PBST and were manually deyolked using forceps. Animal caps were blocked for 1 hour using sheep serum and bovine serum albumin and then incubated in 1:250 anti-β-catenin antibody (BD Bioscience cat# 610154) in blocking solution overnight at 4°C. Animal caps were washed in PBST + 2% DMSO and incubated in 1:200 488 anti mouse antibody (Donkey anti mouse 488, A21202, Life technology) in blocking solution for 2 hours at room temperature. Animal caps were mounted on cover slips and imaged using the Diskovery spinning disc microscope.

### Mass spectrometry

30 embryos at 30% epiboly were dechorionated and placed in a deyolking buffer (55mM NaCl, 1.8mM KCl, 1.25mM NaHCO3). Embryos were vortexed on the lowest speed for 5 minutes to dissolve the yolk and centrifuged at 300 x G to pellet cells. The supernatant was removed, and the cells were washed four times using a wash buffer (110mM NaCl, 3.5mM KCl, 10mM Tris-HCl pH 8.0) followed by 300g spin to pellet cells. Cells were then washed using 1X wash buffer with added EDTA free SIGMAFAST protease inhibitor cocktail tablets. Cells were centrifuged at 300 x G and the final supernatant removed. 500µL of lysis buffer (8mM Urea, 75mM NaCl, 2mM MgCl2, 10mL 1M Tris-HCl pH 8.2, 1X SIGMAFAST protease inhibitor cocktail tablets EDTA free) was added to the cells and sonicated for 10 minutes. Lysates were incubated at 4°C for 30 minutes and centrifuged at 13,000 x G. The supernatant was moved to a clean tube and the pellet was discarded. 25uL of 200mM of iodoacetamide was added to the tubes and incubated for 30 minutes at room temperature in the dark. 1 mL of cold acetone was added, vortexed, and stored at -80°C to precipitate the protein. Protein pellets were then decanted, air dried, and resuspended in 50mM of ammonium bicarbonate. Samples were digested using trypsin and incubated at 37°C overnight and dried using vacuum centrifugation. To concentrate the samples, C18 columns were used, and protein samples were loaded into the Thermo East nLC Orbitrap Exploris 240. Analysis was performed using PEAKS 10 software. Mass spectrometry was performed by the Advanced Analysis Centre at the University of Guelph.

## Authors contributions

IB and TVR conceived and planned the experiments. HK and NS made the *nkd1*^-/-^ mutant. MH, EC, and IB conducted the BIO experiments. HK and IB carried out the qRT-PCR analysis. TH and IB carried out the western blotting analysis. VR and LH carried out the DTU analysis. WL performed Axin2 WMISH and *gsc* arc angle with Axin2 OE and Axin2 MO knockdown. All other experiments were carried out by IB. IB and TVR wrote the manuscript; MH, IB, and TVR edited the manuscript.

## Acknowledgements

We would like to thank Matt Cornish and Mike Davies at the Hagen Aqualab facility at the University of Guelph for supporting the maintenance of the zebrafish system. We would also like to thank Amber Park and Dyanne Brewer at the Advanced Analysis Centre at the University of Guelph for their support in mass spectrometry. Lastly, we would like to thank all current and previous members in the Van Raay lab for all their support.

## Availability of data and materials

Data generated or analyzed during this study are included in this published article [and its supplementary information files]. Complete datasets generated for RNA sequencing during and/or analyzed during the current study are available in the NCBI Gene Expression Omnibus, accession: GSE246858. To review GEO accession GSE246858: Go to https://www.ncbi.nlm.nih.gov/geo/query/acc.cgi?acc=GSE246858.

