## Supplemental figures for "Loss of Nkd1 is epistatic to loss of Axin2 in regulating Wnt signaling"

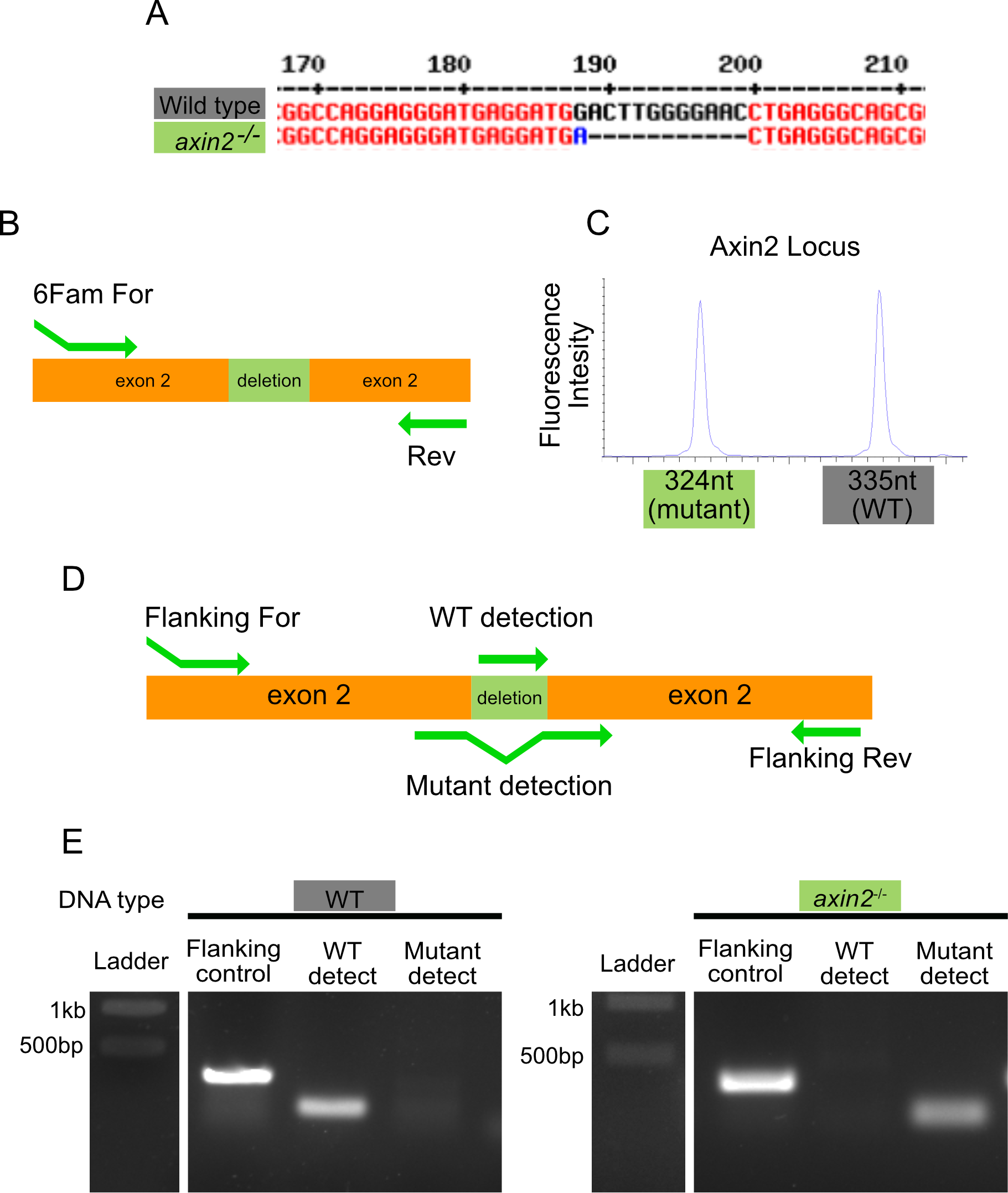

**Supplemental figure 1: Confirming the Axin2 mutant allele.** (A) The Axin2 mutant allele in the founder *axin2*^-/-^ zebrafish was sequenced and had a 12nt deletion (shown in black) with a 1 nucleotide insertion (shown in blue). (B, C) Fragment analysis was used to detect amplicon size of the region with the mutant allele being 11nt smaller than the wild type allele. (D, E) A PCR based genotyping approach was developed for the Axin2 mutant allele.

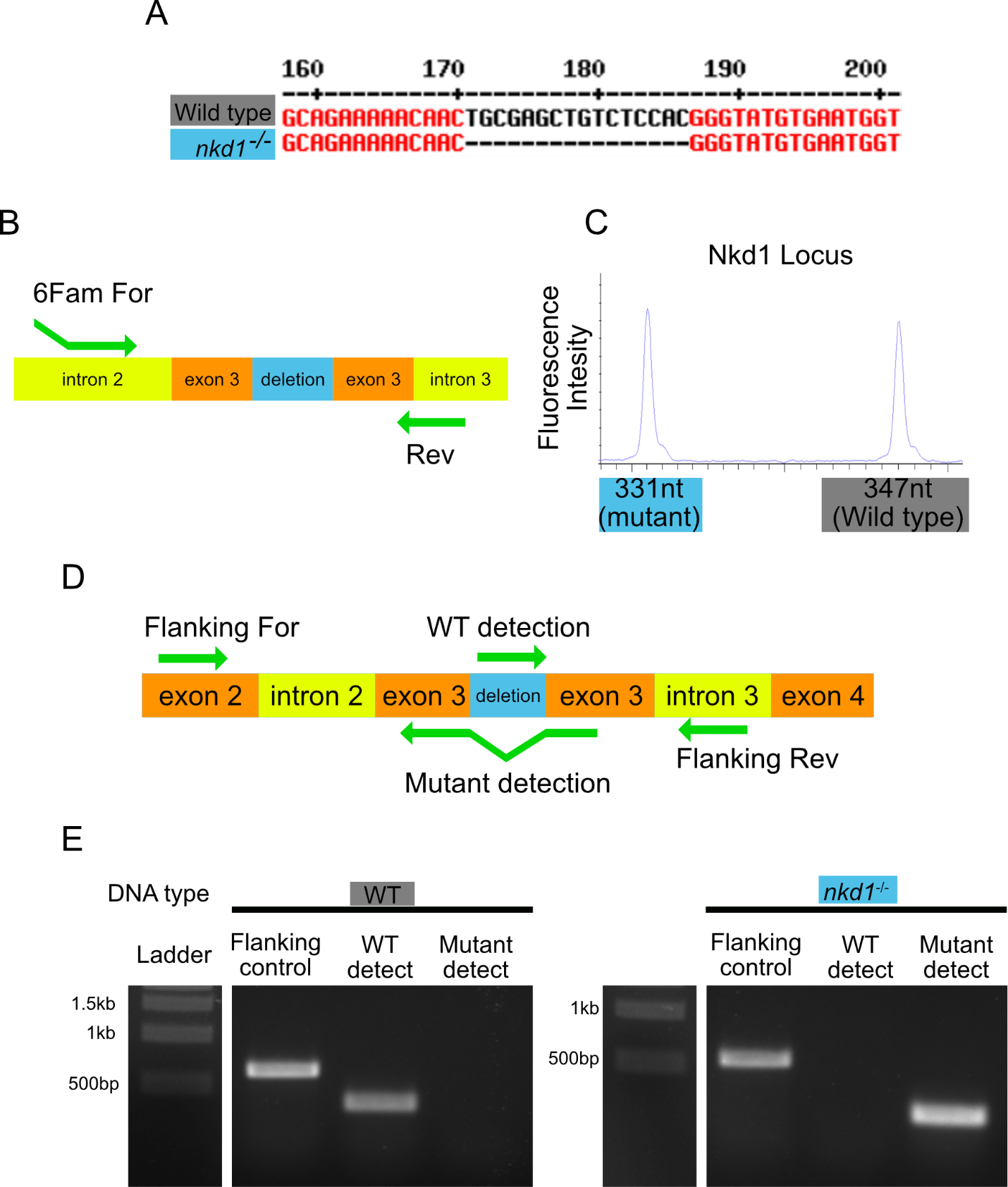

**Supplemental figure 2: Confirming the Nkd1 mutant allele**. (A) The Nkd1 mutant allele in the founder *nkd1*^-/-^ zebrafish was sequenced had a 16nt deletion (shown in black). (B, C) Fragment analysis was used to detect amplicon size of the region with the mutant allele being 16nt smaller than the wild type allele. (D, E) A PCR based genotyping approach was developed for the Nkd1 mutant allele.

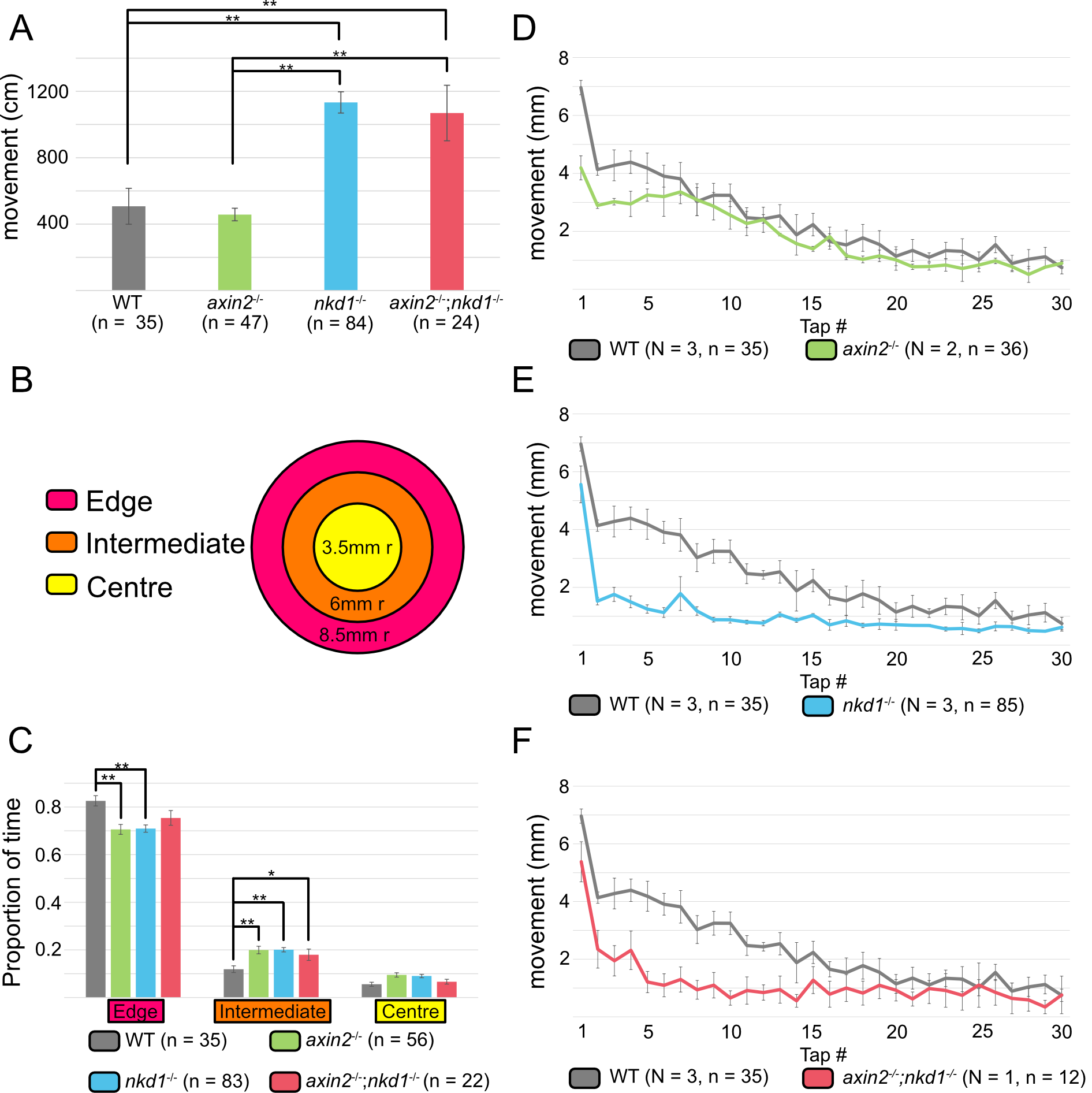

**Supplemental figure 3: Wnt regulator mutants have distinct swimming behavior and reduced startle response**. (A) At 5dpf zebrafish larvae were recorded for 1.5 hours in the dark with *nkd1*^-/-^ and *axin2*^-/-^;*nkd1*^-/-^ having increased total movement. (B-C) All the Wnt regulator mutants had an increased proportion of time spent in the intermediate zone of the well when compared to wild type larvae. (D-F) After the 1.5 hour dark incubation, a tap analysis was performed where both *nkd1*^-/-^ and *axin2*^-/-^;*nkd1*^-/-^ had reduced startle response. A one-way ANOVA was used for all statistics with p-value < 0.05 = *, p-value < 0.01 = **. Behavior analysis was conducted in a 24 well plate using DanioVision temperature control unit (Noldus Information Technology) and video recordings were analyzed using EthoVision XT 16.

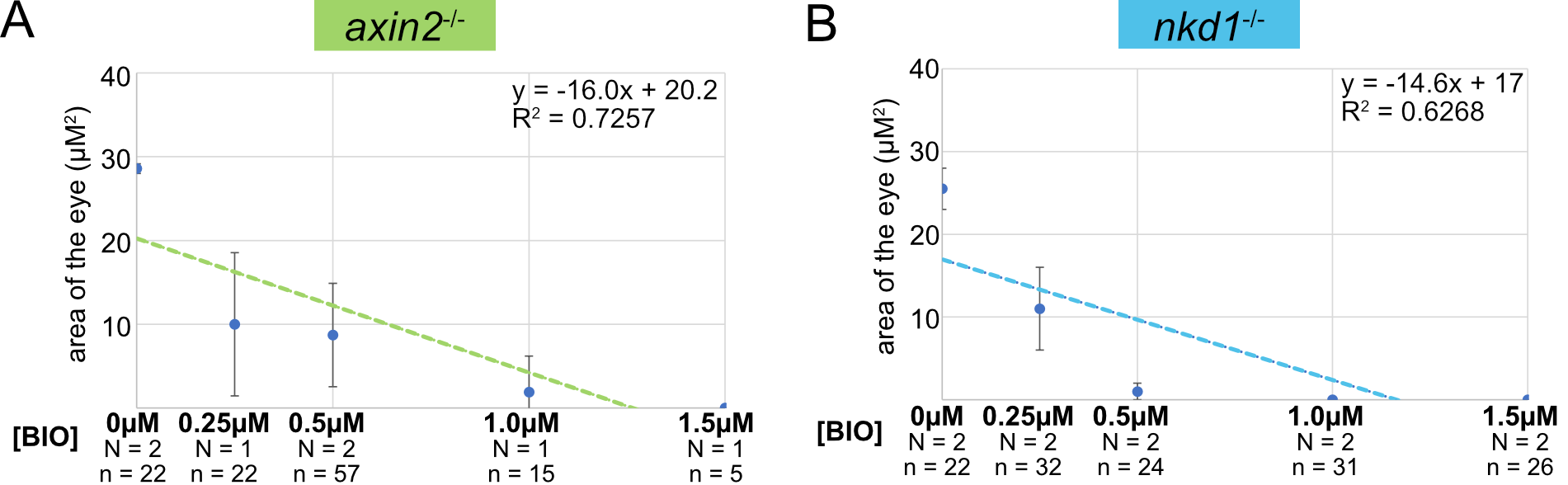

**Supplemental figure 4. The *axin2*^-/-^ and *nkd1*^-/-^ mutants are sensitive to BIO.**

(A) *axin2*^-/-^ and (B) *nkd1*^-/-^ embryos were treated with BIO from 4 – 30 hpf and measurements were taken on ImageJ. Error bars represent SEM when N = 2. Error bars represent standard deviation when N = 1.

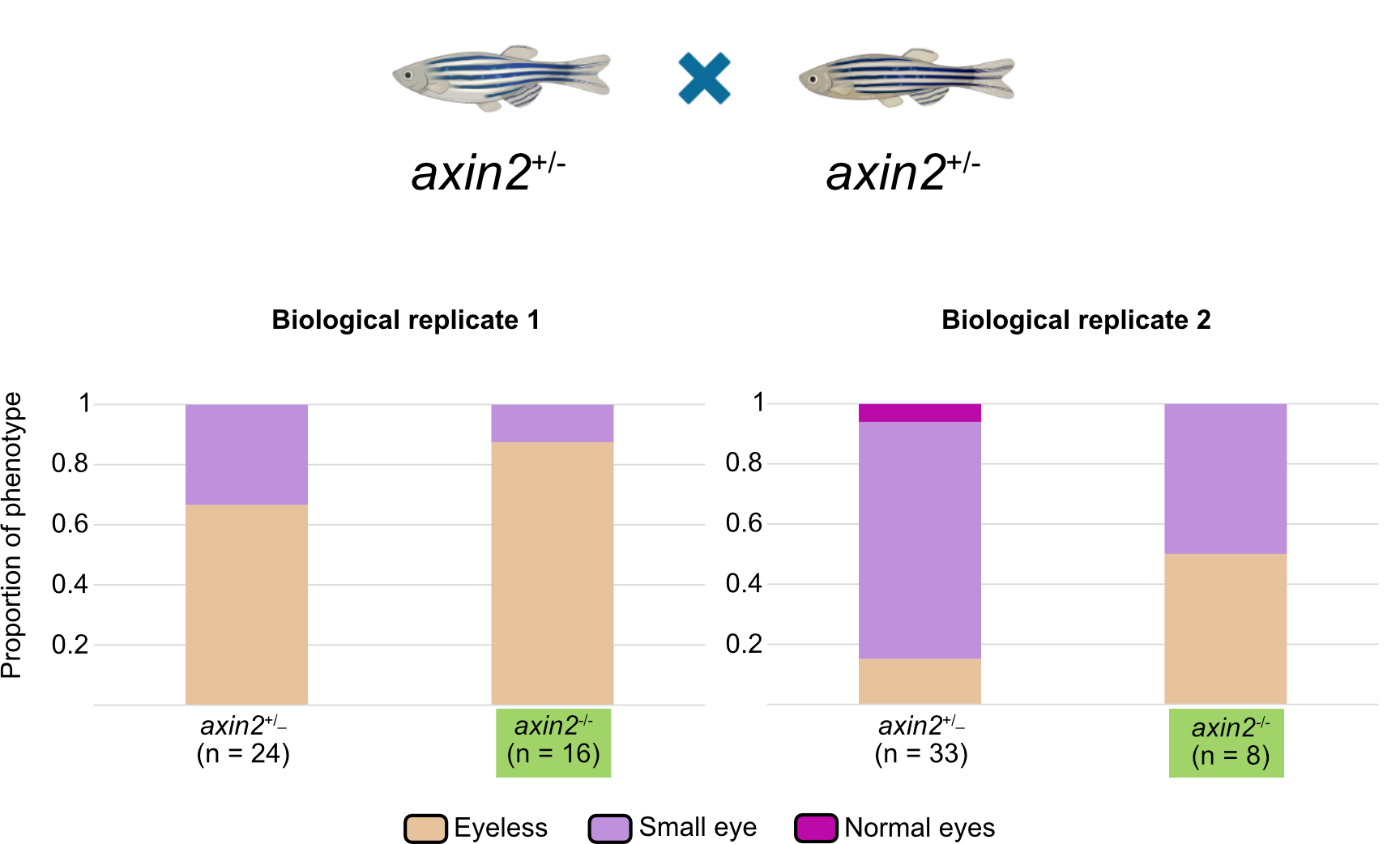

**Supplemental figure 5. *axin2*^-/-^ mutants are sensitive to BIO.**

Embryos from an axin2^+/-^ incross were treated with 0.75µM of BIO + 2% DMSO from dome stage to 1dpf. Phenotypes were recorded at 1dpf and was seen that the *axin2*^-/-^ embryos had an increase in eyeless phenotype when compared to *axin2*^+/_^ embryos from the same cross.

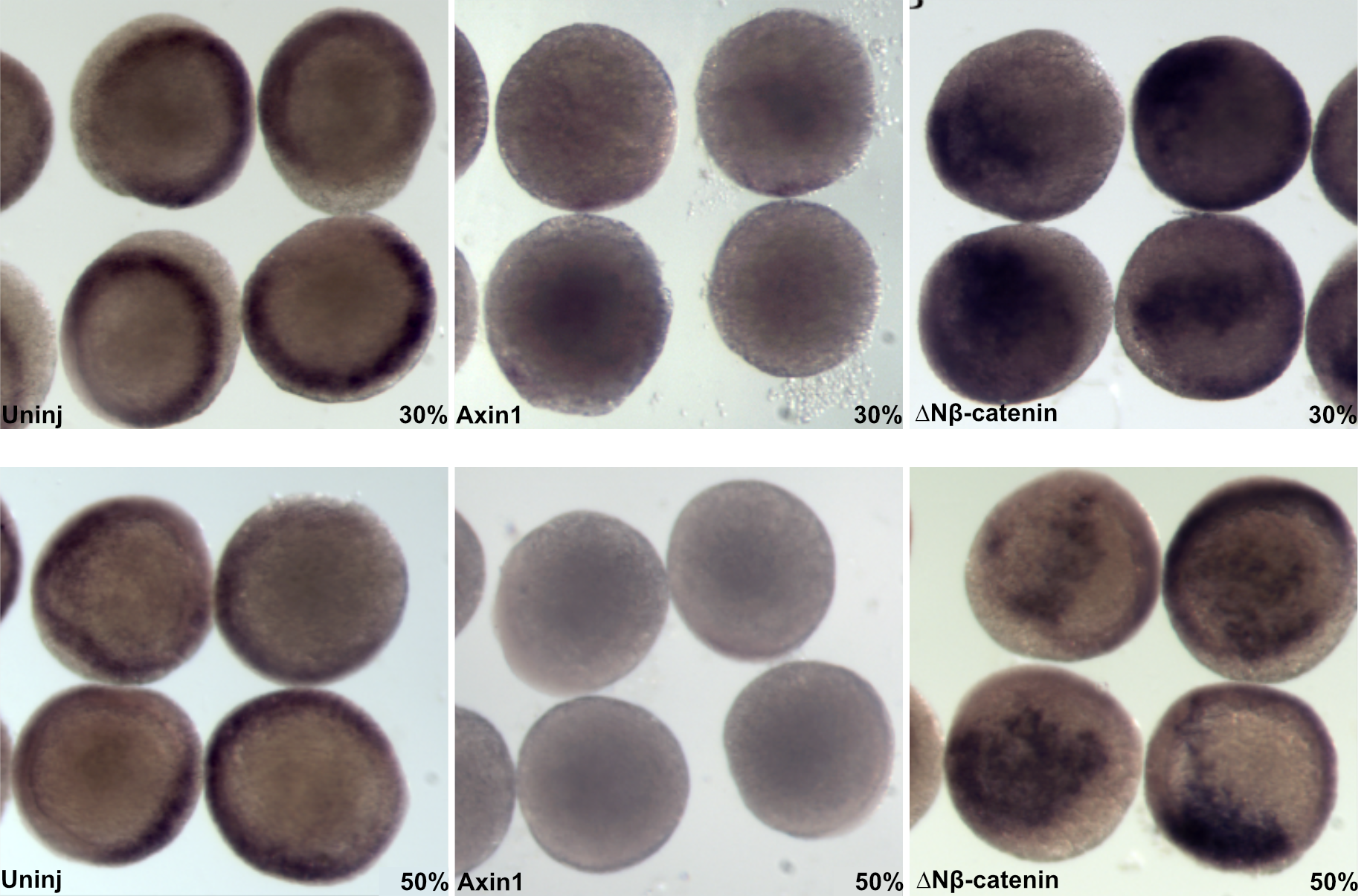

**Supplemental figure 6. Wnt signaling is necessary and sufficient for *axin2* expression at 30% and 50% epiboly.** WMISH using an *axin2* anti-sense probe was used to assess *axin2* expression with hyperactivation or hypoactivation of Wnt signaling at either 30% or 50% epiboly with images taken in an animal view. Injections were performed at the one cell stage with either *axin1* or *ΔβN-catenin*. At both 30% and 50% epiboly *axin2* expression is restricted to the ventrolateral domain. Inhibiting Wnt signaling by overexpressing *axin1* abolished *axin2* expression, whereas activated the pathway with overexpression of *ΔNβ-catenin* caused ectopic expression of *axin2* at both 30% and 50% epiboly. (N = 2)

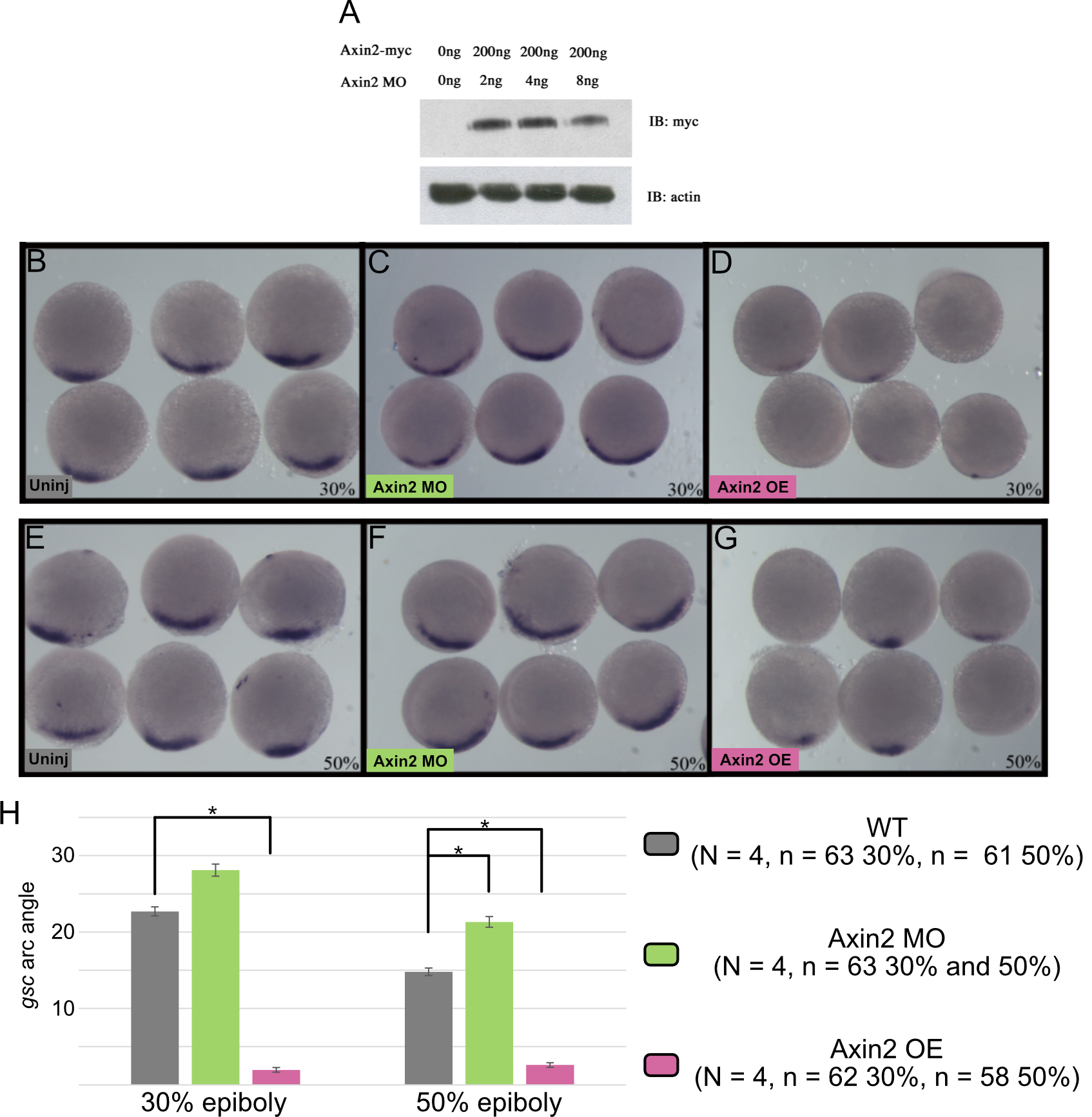

**Supplemental figure 7. Knocking down Axin2 reduces *gsc* arc at 50% epiboly.** (A) verification of Axin2 morpholino (MO) efficacy following co-injection with 2-8ng of Axin2 MO, western blot analysis of 10 embryos (pooled) revealed a decrease in Axin2 expression as MO concentration increased. (B-D) The arc of *gsc* was not affected by knocking down Axin2 using MO but was severely reduced with Axin2 overexpression at 30% epiboly. (E-G) The arc of *gsc* was increased by knocking down Axin2 using MO and was severely reduced with Axin2 overexpression at 50% epiboly. (E) The *gsc* arc was measured using an eye piece attached to the microscope (arc angles of *gsc* are vastly different than Figure 5 due to the change in how it was measured, however a similar trend was found). Error bars represent standard error (Bonferroni post-hoc, *, p< 0.05).

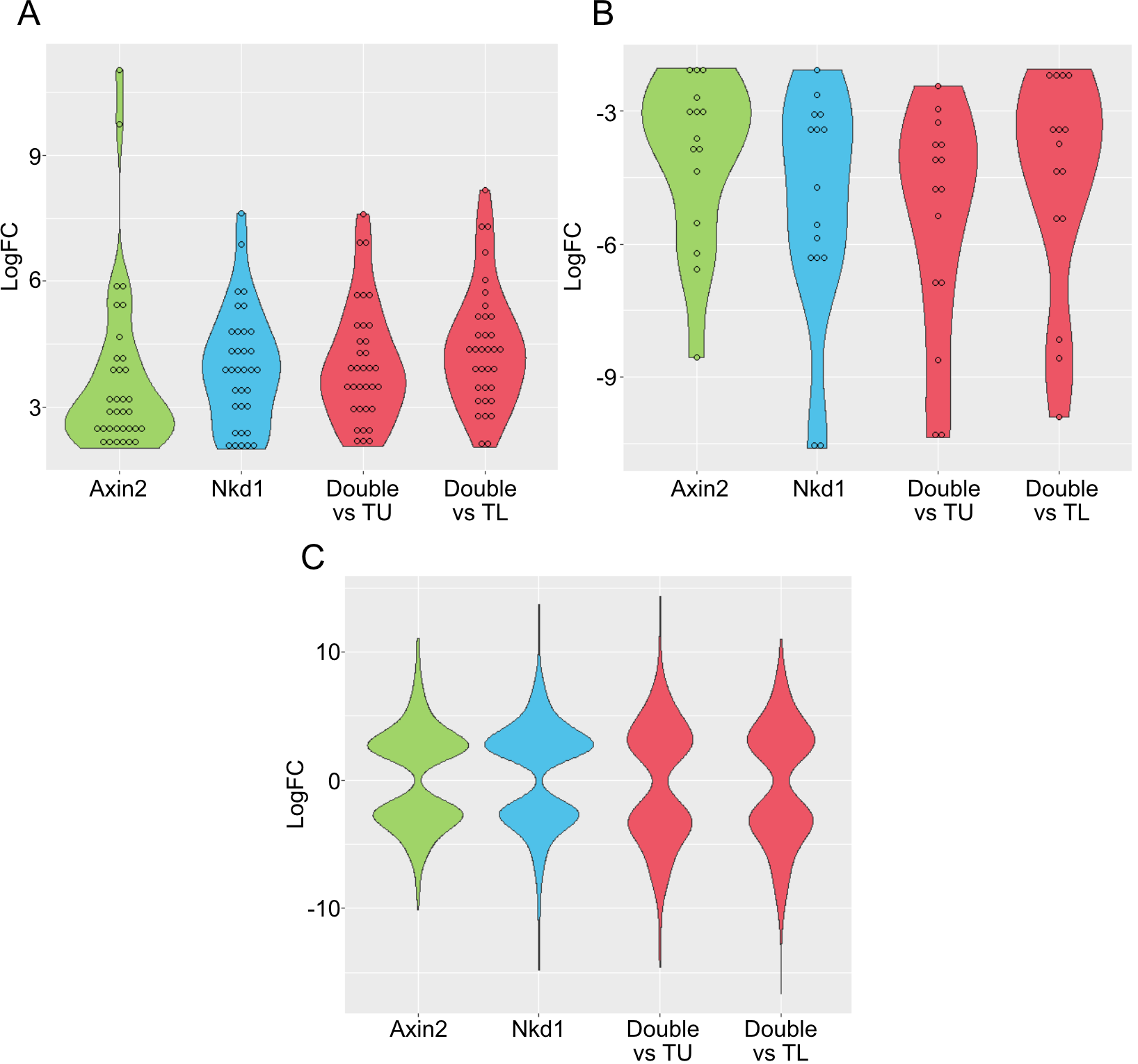

**Supplemental figure 8. Violin plots for RNA-seq data.** (A) 34 Up-regulated genes with a logFC > 2 and FDR < 0.05 for all Wnt regulator mutants when compared to respective background genotype. (B) 15 down-regulated genes with logFC < -2 and FDR < 0.05 for all Wnt regulator mutants when compared to respective background genotypes. (C) All differentially expressed genes in each genotype with an absolute logFC > 2 when compared to background genotype. Genes were only included for the *axin2*^-/-^;*nkd1*^-/-^ if they were significant when compared to both TU and TL backgrounds. 735 DEG for *axin2*^-/-^ vs TL, 869 DEG for *nkd1*^-/-^ vs TU, and 312 DEG for *axin2*^-/-^;*nkd1*^-/-^ vs TU and TL.

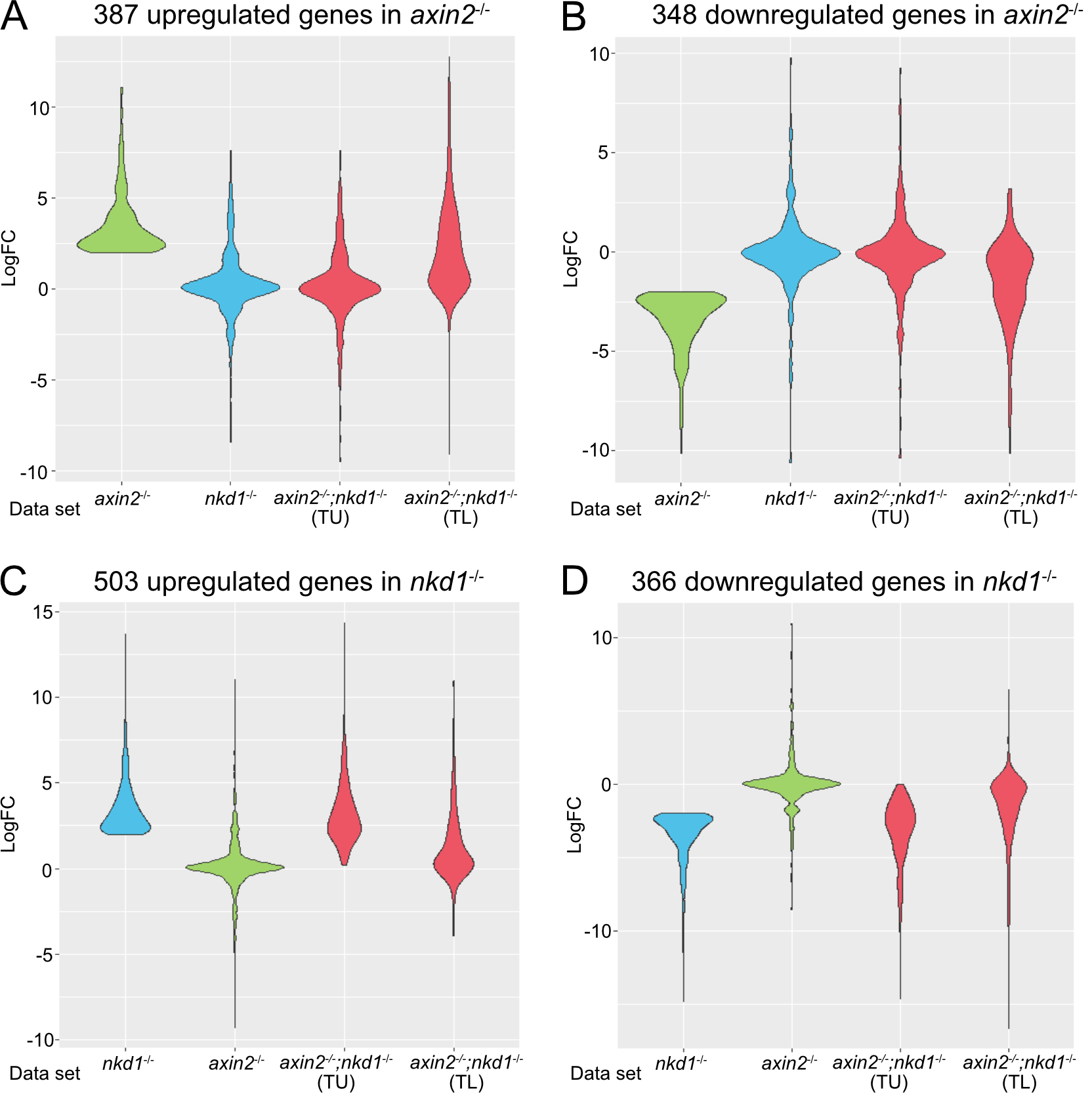

**Supplemental Figure 9. DEG in *axin2*^-/-^ are unchanged in *nkd1*^-/-^ and *axin2*^-/-^;*nkd1*^-/-^**. (A) The 387 upregulated genes (logFC > 2, FDR < 0.05) and (B) 348 downregulated genes (logFC < -2, FDR < 0.05) in *axin2*^-/-^ were used to evaluate the raw expression values for *nkd1*^-/-^ and *axin2*^-/-^;*nkd1*^-/-^. (C) The 503 upregulated genes (logFC > 2, FDR < 0.05) and (D) 366 downregulated genes (logFC < -2, FDR < 0.05) in *nkd1*^-/-^ were used to evaluate the raw expression values for *axin2*^-/-^ and *axin2*^-/-^;*nkd1*^-/-^.

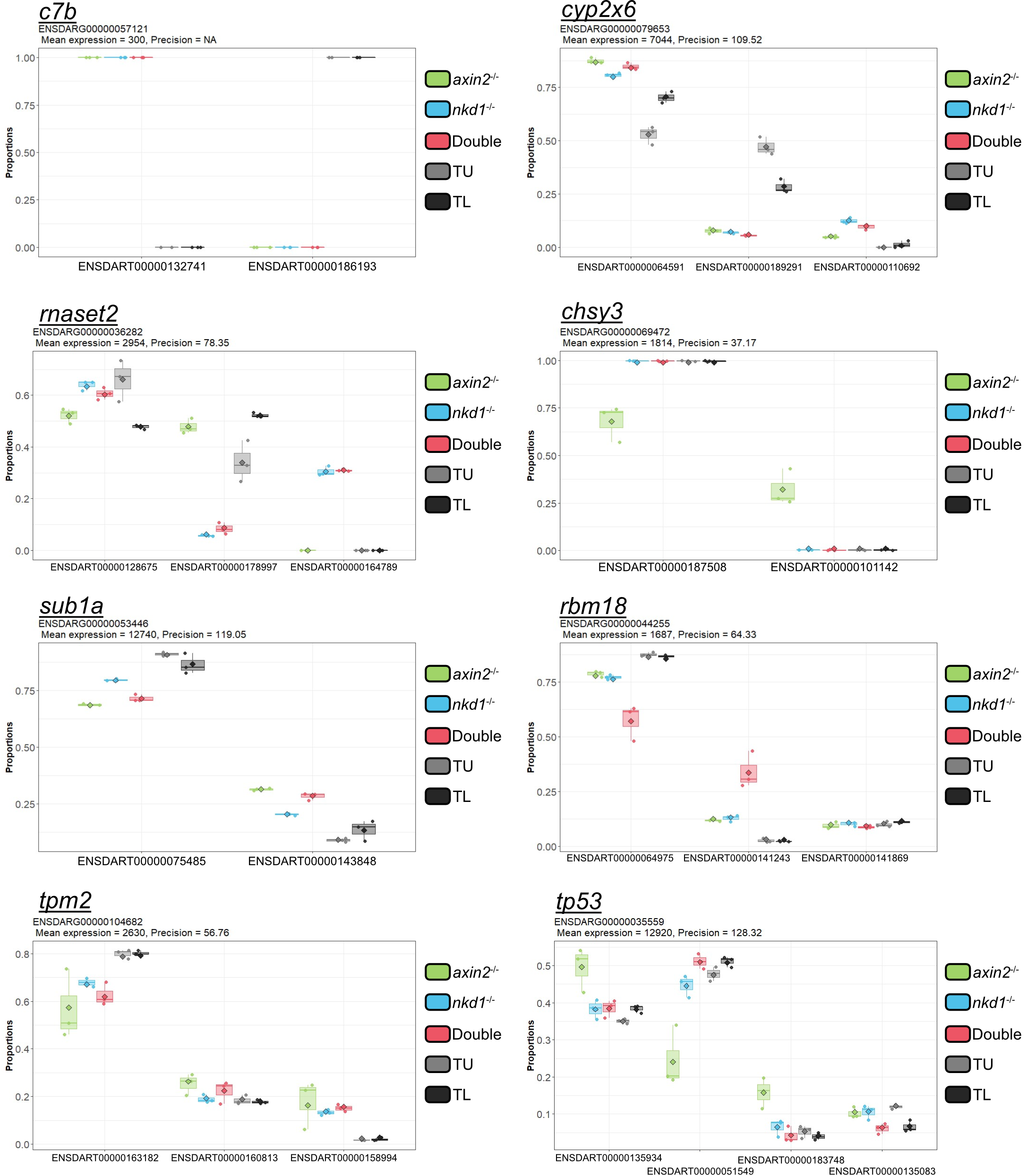

**Supplemental figure 10: Top hits differential transcript usage #1.**

Top hits were determined by having the smallest p-value in the analysis as well as having a similar DTU between the TL and TU groups. DTU analysis was performed using DRIMSeq.

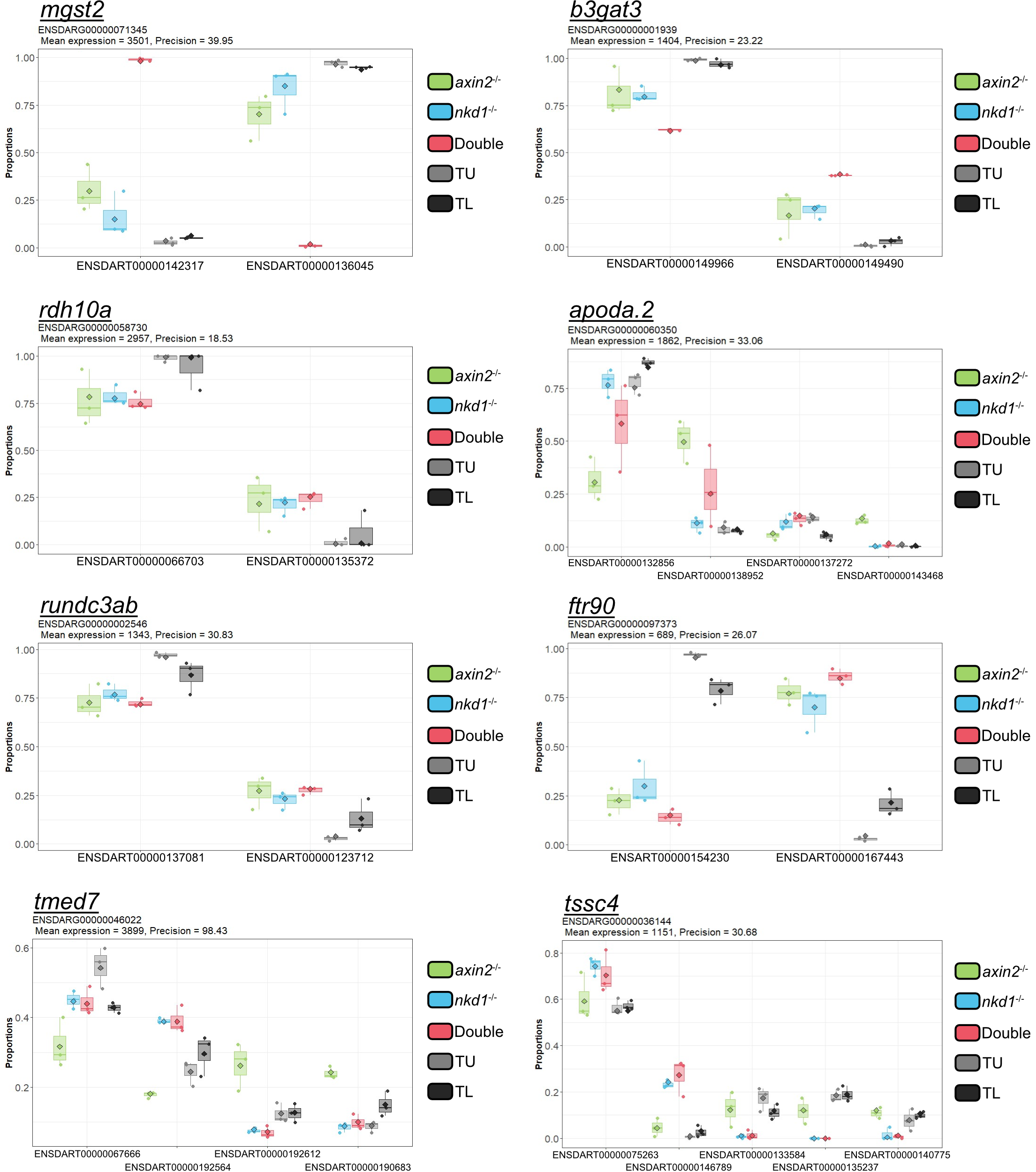

**Supplemental figure 10 continued: Top hits differential transcript usage #2**.

Top hits were determined by having the smallest p-value in the analysis as well as having a similar DTU between the TL and TU groups. DTU analysis was performed using DRIMSeq.

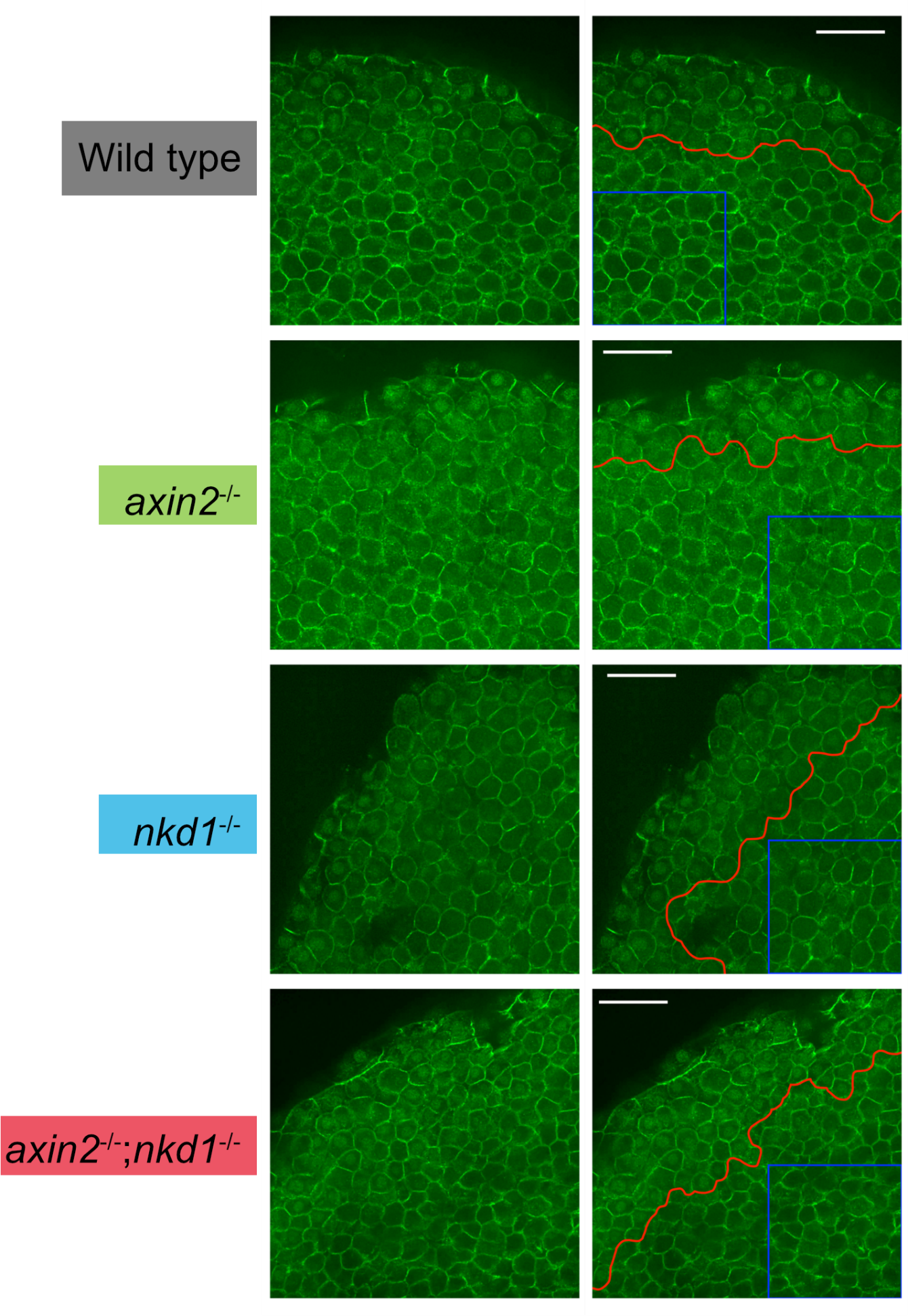
**Supplemental figure 11. Wnt regulator mutants have no change in nuclear β-catenin levels at 30% epiboly.** Immunohistochemistry was performed at 30% epiboly for β-catenin with images taken at the edge of the embryos to capture the ventrolateral domain which has endogenous Wnt signaling (marked in red). Cells that were not in the ventrolateral border were assessed for nuclear β-catenin accumulation with none of the Wnt regulator mutants showing any sign of increased nuclear β-catenin (blue box). Scale bars = 50µM
